# Single-step genome engineering in the bee gut symbiont *Snodgrassella alvi*

**DOI:** 10.1101/2023.09.19.558440

**Authors:** Patrick J. Lariviere, A. H. M. Zuberi Ashraf, Sean P. Leonard, Laurel G. Miller, Nancy A. Moran, Jeffrey E. Barrick

**Affiliations:** Department of Molecular Biosciences, The University of Texas at Austin, Austin, TX 78712, USA; Department of Integrative Biology, The University of Texas at Austin, Austin, TX 78712, USA

## Abstract

Honey bees are economically relevant pollinators experiencing population declines due to a number of threats. As in humans, the health of bees is influenced by their microbiome. The bacterium *Snodgrassella alvi* is a key member of the bee gut microbiome and has a role in excluding pathogens. Despite this importance, there are not currently any easy-to-use methods for modifying the *S. alvi* chromosome to study its genetics. To solve this problem, we developed a one-step procedure that uses electroporation and homologous recombination, which we term SnODIFY (**Sn**odgrassella-specific **O**ne-step gene **D**eletion or **I**nsertion to alter **F**unctionalit**Y**). We used SnODIFY to create seven single-gene knockout mutants and recovered mutants for all constructs tested. Nearly all transformants had the designed genome modifications, indicating that SnODIFY is highly accurate. Mutant phenotypes were validated through knockout of Type 4 pilus genes, which led to reduced biofilm formation. We also used SnODIFY to insert heterologous sequences into the genome by integrating fluorescent protein-coding genes. Finally, we confirmed that genome modification is dependent on *S. alvi*’s endogenous RecA protein. Because it does not require expression of exogenous recombination machinery, SnODIFY is a straightforward, accurate, and lightweight method for genome editing in *S. alvi*. This workflow can be used to study the functions of *S. alvi* genes and to engineer this symbiont for applications including protection of honey bee health.

## Introduction

Honey bees are critical pollinators in agriculture and are essential for global food security^1–3^. Both managed and wild bee populations are experiencing population declines due to a number of stressors that include viral and bacterial pathogens^4–6^. It has become increasingly clear that the health of bees is influenced by their microbiome^7–12^. One of the eight core gut microbiome constituents is *Snodgrassella alvi* (*Neisseriaceae*) which colonizes the wall of the hindgut ileum^7,13^. *S. alvi* is a keystone species within the gut community, as it depletes oxygen and thereby creates an anaerobic environment for other community members^9^.

Experimental studies with gnotobiotic bees also show that *S. alvi* also protects against gut pathogens^14–16^. Further, *S. alvi* containing engineered plasmids have been used to induce RNAi to perform functional genomics in bees^17,18^, and to combat bee pathogens such as *Nosema* and deformed wing virus^16,17,19^. However, fundamental questions still remain concerning how *S. alvi* colonizes and interacts with its bee host.

To date, genetic studies of *S. alvi* have primarily been limited to -omics level investigations employing Tn-seq and RNA-seq^7,14,20,21^. Additionally, applications aimed at studying and improving bee health have been constrained to the use of wild type or plasmid-bearing *S. alvi*^16–18,22,23^. As a result, research involving *S. alvi* has been hampered by the lack of streamlined tools for reverse genetics. Gene disruption in *S. alvi* was previously achieved through conjugation of a suicide vector containing an antibiotic resistance gene flanked by homology to the chromosome and a small guide RNA targeting cleavage of the integration site by Cas9 expressed from a second plasmid^22^. However, this conjugation-dependent method is technically complex and can lead to off-target single-crossover integrations, necessitating several selection and screening steps to isolate an on-target knockout. Therefore, there is a need for a more accurate and straightforward genome engineering workflow in *S. alvi*.

Homologous recombination-based gene knockout^24–28^ has long been employed for genome modification in bacteria. For this approach to be successful, one must deliver a sufficient quantity of DNA into cells, and cells must have native or exogenous recombination pathways that are efficient enough to reliably integrate this DNA into their genomes. Recombination with the native RecA pathway has been used for modifying the chromosomes of several species of *Neisseria*, which are close relatives of *Snodgrassella* species^29^. In these cases, delivery of the knockout DNA has typically relied on natural transformation^30–32^. *S. alvi*, however, appears to not be naturally competent. Its genome does not contain repeated DNA uptake-sequences characteristic of *Neisseria* species^33^, and it does not transform DNA in normal laboratory culture conditions. This has limited prior work using homologous recombination for genome modification to the delivery of DNA through conjugation, which is not as amenable to making fast and accurate genome modifications as is transforming linear double-stranded DNA fragments.

We have overcome the DNA delivery hurdle by demonstrating that *S. alvi* can be transformed by electroporation and find that *S. alvi*’s native RecA pathway is efficient enough to reliably use homologous recombination for genome modification. Here, we describe a streamlined, lightweight, and highly accurate method to modify the *S. alvi* genome, which we call SnODIFY (**Sn**odgrassella-specific **O**ne-step gene **D**eletion or **I**nsertion to alter **F**unctionalit**Y**). SnODIFY will enable future studies of *S.alvi*-host interactions and applications that engineer this symbiont to protect bee health. Beyond *S. alvi*, the principles underlying this workflow may inform genome engineering technology development in other insect symbionts.

## Results

### *S. alvi* can be transformed by electroporation

To determine if *S. alvi* could be transformed by electroporation, we prepared electrocompetent *S. alvi* wkB2^29^ cells in a manner similar to that used for *Escherichia coli*^34^. Cells grown in cell culture flasks containing Columbia broth were washed twice with dH_2_O, then resuspended in 10% glycerol. Electrocompetent cells were electroporated with 520 ng of the *specR-*containing plasmid pBTK800, which is capable of replicating in *S. alvi* and has previously been transferred to *S. alvi* by conjugation^18^. Transformed cells were recovered overnight, and plated on selective media. Cells that had received pBTK800 were able to grow on plates containing spectinomycin, whereas those that received no DNA could not (Fig. S1), indicating that growth is not due to spontaneous mutations in the chromosome that can yield spectinomycin resistance in other bacteria. These results demonstrate that *S. alvi* can be transformed with plasmids by electroporation.

### Knockout of *staA* by transforming dsDNA

We next investigated whether electroporation of linear double-stranded DNA (dsDNA) could be leveraged to modify the *S. alvi* chromosome. We devised a strategy, SnODIFY, to knockout a gene of interest by homologous recombination (Fig. 1A). In the SnODIFY workflow, a construct is designed with homology arms to a gene of interest, flanking an antibiotic resistance cassette. Linear DNA constructs are chemically synthesized, assembled, and PCR amplified. Then, *S. alvi* wkB2 cells are electroporated with the dsDNA, recovered overnight, selected on antibiotic-containing media, and screened for proper insertion of the construct (Fig. 1B).

**Fig 1.**
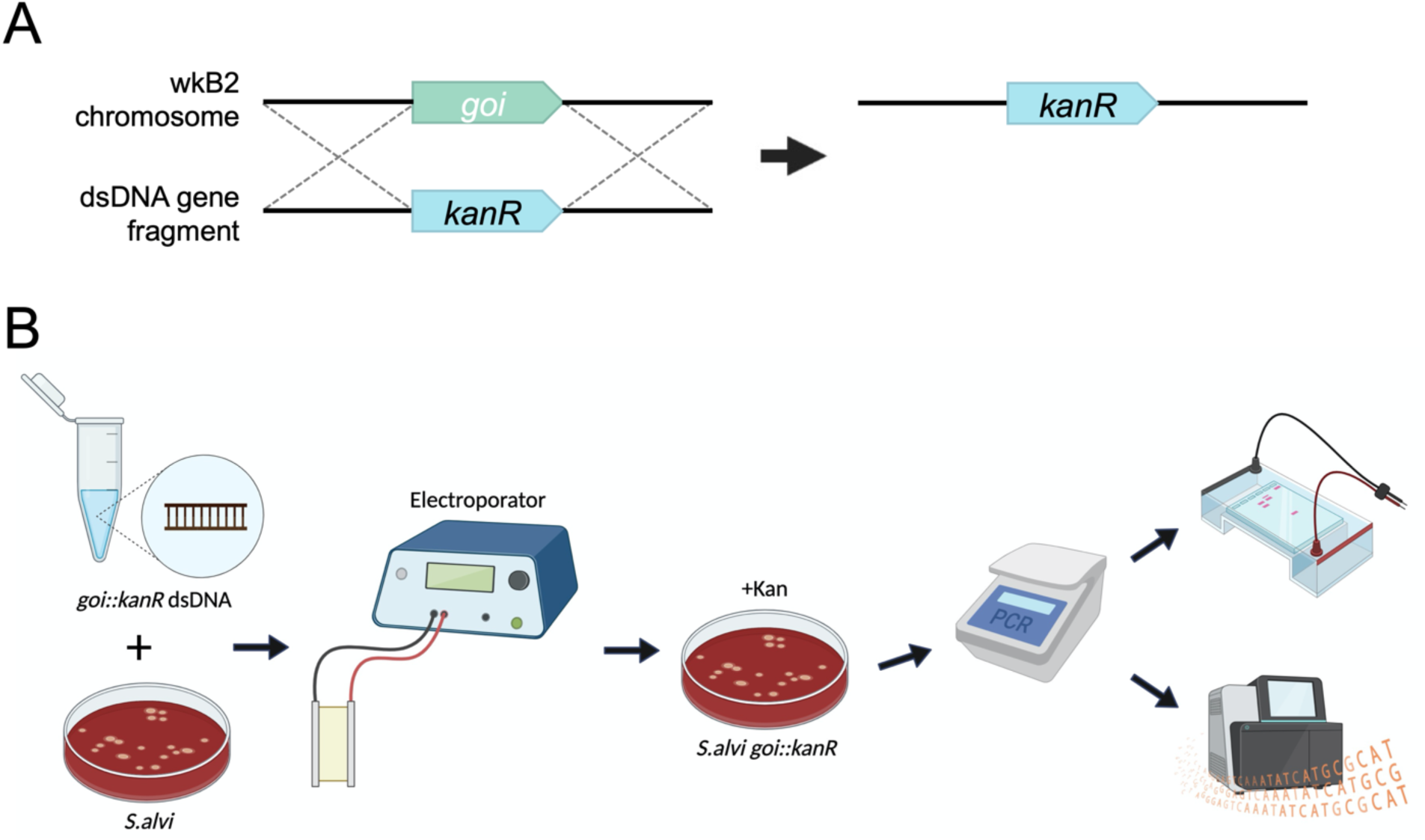
SnODIFY overview and workflow. A. Cartoon depicting an overview of gene knockout by SnODIFY. An antibiotic resistance cassette, such as *kanR*, is designed to be flanked by arms with homology surrounding the deletion target. The targeted genomic locus is replaced by the antibiotic resistance cassette by single-step homologous recombination. B. Diagram of SnODIFY workflow. 5 μg of linear or circular knockout construct DNA is electroporated into *S. alvi* wkB2. Following overnight liquid recovery with no antibiotics, transformants are selected for on media containing antibiotic. Transformants are isolated and the deleted region is screened via PCR. Amplicons are purified and visualized on a 1% agarose gel, then sent for amplicon/whole-genome sequencing and aligned to a reference sequence.

To pilot this method, we targeted *staA* for deletion. This gene is non-essential^21^ and was previously disrupted using the Cas9/suicide vector approach^22^. We designed *kanR* knockout constructs with homology arms inside of the *staA* gene that were either 500 or 1000 bp long (Fig 2A). Initial electroporations using approximately 150 ng and 300 ng of DNA did not yield any transformants. Increasing the concentration of DNA to 5 μg proved sufficient for isolating kanamycin resistant transformants (Fig 2B). We found that 1000 bp of homology yielded 85 transformants, whereas 500 bp of homology yielded none (Fig 2B; Table S1). From the plate transformed with 1000 bp homology arms, we picked seven large and seven small colonies. Colonies were screened via PCR for insertion of the *kanR* cassette using primers specific to sequences within the homology arms. All 14 of the screened colonies contained the *kanR* cassette inserted at the proper locus (Fig 2C; Table S1), indicating on-target editing accuracy of 100% in the screened colonies. Short-read whole-genome sequencing (WGS) of one of these *staA::kanR* strains confirmed that the targeted stretch of *staA* had been deleted and replaced with the *kanR* gene (Table S2).

**Fig 2.**
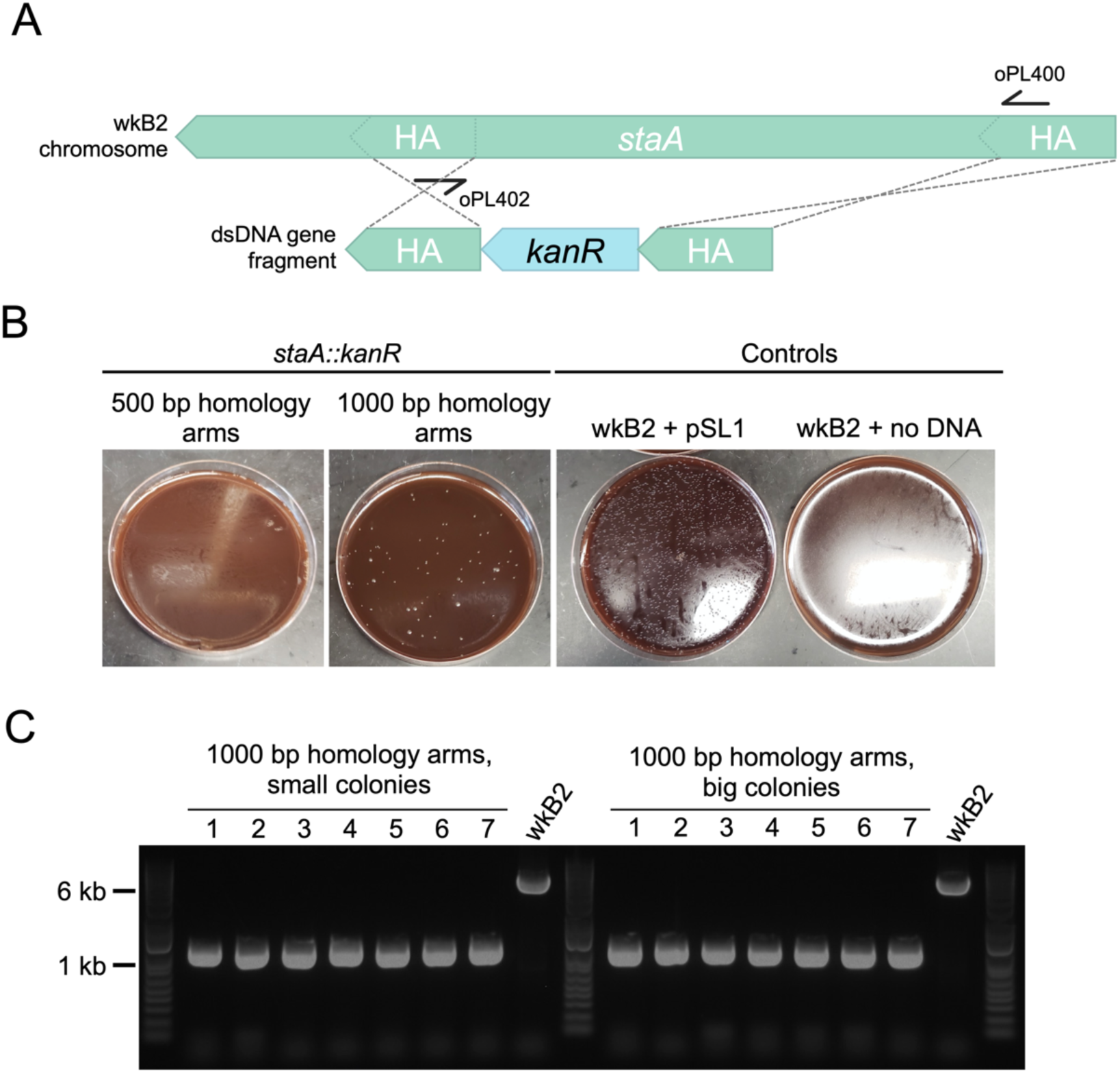
Gene knockout in *staA* is highly accurate. A. Gene deletion map depicting genomic *staA* and the corresponding deletion construct. The deletion construct is composed of a *kanR* cassette flanked by 500 or 1000 bp arms with homology internal to the *staA* gene. Primers oPL400 and oPL402 were designed for use in PCR screening for proper deletion. B. Images of *staA::kanR* Columbia + 5% sheep’s blood (Col-B) agar plates + kanamycin transformation plates. wkB2 can be successfully transformed with a *staA::kanR* cassette containing 1000 bp (middle left), but not 500 bp (left), homology arms, indicating that 1000 bp of homology is needed to isolate colonies with the deletion. wkB2 cannot grow on kanamycin plates (right) without the presence of a kanamycin resistance cassette, as illustrated by control cells that received the *kanR*-containing pSL1 plasmid (middle right). C. DNA gel of PCR screen for *staA* deletion. Large and small colonies of wkB2 + *staA::kanR* with 1000 bp homology arms or wkB2 were amplified via PCR using oPL400 and oPL402, and run on a 1% agarose gel. The ∼6000 bp band observed in the wkB2 sample corresponds to the genomic region within *staA*, whereas the ∼1000 bp band observed in experimental samples corresponds with the *kanR* cassette. 100% of both small and large colonies were found to contain a single 1000 bp band, indicating proper deletion of *staA*, regardless of colony size.

### Genes can be reliably knocked out using SnODIFY

To further assess the reliability of SnODIFY, we designed and tested knockout constructs for six additional non-essential^21^ *S.alvi* genes: *pilF*, *pilG*, *asr*, *asmE*, *recJ*, and *recA*. Similar to the *staA* knockout construct, we designed kanamycin resistance cassettes flanked by 1000 bp homology arms (Fig. S2). However, this time, the homology arms were designed to match regions upstream and downstream of the CDS, with 10 bp of CDS left on 5’ and 3’ ends, so that nearly the whole gene would be deleted (Fig. S2). As before, dsDNA constructs were synthesized and electroporated into *S. alvi* wkB2. Then, cells were plated on antibiotic-containing media to select for integration of the cassette into the chromosome. The number of transformants ranged from ∼20 to >100 colonies, depending on the gene targeted for knockout (Table S1). Because editing accuracy was found to be 100% for the *staA* knockout, in some cases, we screened fewer clones for successful deletion by PCR for these six additional knockouts. For five out of the six knockouts, editing accuracy (as determined by PCR or WGS) was 100% (Fig. S3; Table S1). For the remaining gene, *asr*, editing accuracy was 50% (Fig. S3; Table S1). (Inconclusive PCR screening of *asr*, evidenced by the presence of two bands instead of one (Fig. S3), may mask a higher editing accuracy). WGS confirmed each of these six genes was knocked out and successfully replaced by the kanamycin resistance cassette (Table S2). In some cases, point mutations or small (primarily ≤ 20 bp; in one case, 69 bp) deletions distant from the integration site were observed (Table S2; Table S3). In aggregate, these data demonstrate the feasibility of scaling up SnODIFY to construct multiple *S. alvi* knockout strains in parallel, with near-perfect on-target editing accuracy.

### SnODIFY-generated type IV pilus mutants have impaired biofilm formation

Biofilm formation has previously been used as a phenotype for validating novel genome engineering tools^35^. Two of the *S. alvi* knockout strains we created had deletions of type 4 pilus genes, *pilF* and *pilG*. Biofilm formation was quantified for wild type wkB2 and the *pilF::kanR* and *pilG::kanR* mutants using a crystal violet assay, by assessing OD_550_/OD_600_ of washed cells grown in 96 well plates. We found that knockout of both *pilF* and *pilG* led to a significant and visually noticeable decrease in biofilm formation compared to the wild type strain (Fig 3; Fig S4), similar to that observed for mutant strains containing *pilF* and *pilG* genes disrupted by transposon insertions in a prior study^21^. This result indicates SnODIFY can be used to generate knockouts for downstream phenotype testing.

**Fig 3.**
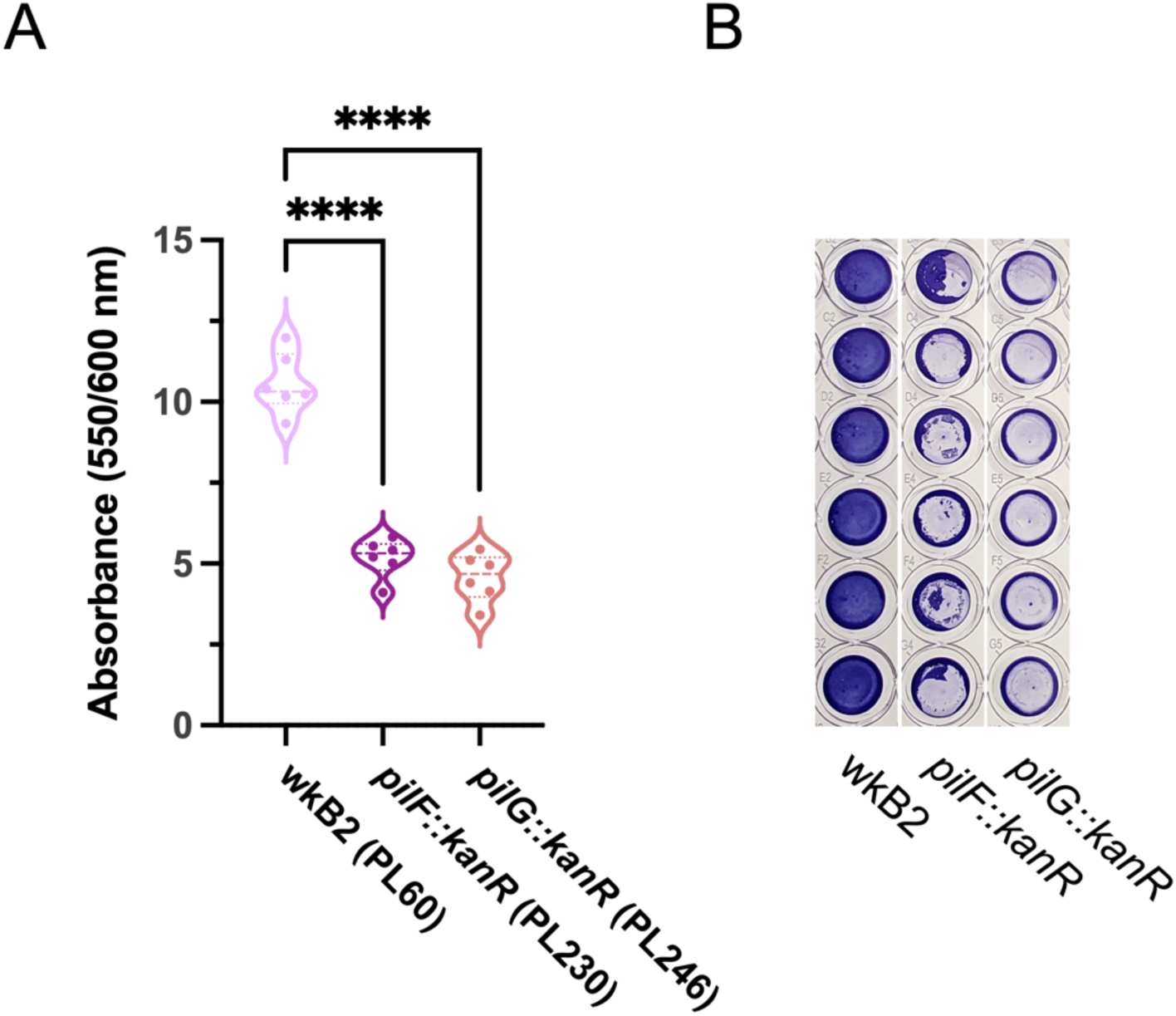
Phenotypic validation: Knockout of genes encoding adhesion proteins result in biofilm formation defect. A. Plot of OD_550_/OD_600_ to quantify biofilm formation normalized to cell growth for biofilm knockout strains. Compared to WT, *pilF::kanR* and *pilG::kanR* have decreased biofilm formation relative to growth. Thick dotted line = median; thin dotted line = quartiles. N=6 for each condition. ****: P < 0.0001, as determined by One-way parametric ANOVA with Dunnett’s multiple comparison test. B. Image of Crystal violet-stained 96 well plate columns demonstrating *pilF::kanR* and *pilG::kanR* mutants have reduced biofilm formation as compared to wkB2.

### Heterologous genomic insertions can be made using circular plasmid DNA

We next sought to implement two additional features of SnODIFY: insertion of heterologous genes into the genome; and genome modification using circular DNA, as opposed to linear DNA. We approached these problems simultaneously through insertion of fluorescent protein-coding genes into the chromosome using electroporation of *E. coli* plasmids that do not replicate in *S. alvi*. We reasoned that recombination using plasmid DNA would allow for knockout cassettes to be constructed from part plasmids according to the bee microbiome toolkit (BTK) assembly scheme^22^, which would enable re-use of the same homology flanks to easily integrate different payloads into the genome.

To pilot this approach, we identified an intergenic region of the wkB2 genome, situated between the non-essential genes *SALWKB2_RS11215* and *SALWKB2_RS11220*^21^. We engineered part plasmids containing genes encoding one of two fluorescent proteins (GFP or E2-Crimson) and an antibiotic resistance gene, as well as homology arm part plasmids (Fig 4A, B; Fig S5; Fig S6). Parts were assembled into a single plasmid via Golden Gate cloning in *E. coli*^22^. The purified non-linearized plasmid was then electroporated into *S. alvi* followed by isolation of antibiotic resistant colonies (Fig S5; Fig S6). Insertion of the cassette into the proper genomic location was confirmed by WGS (Table S2).

**Fig 4.**
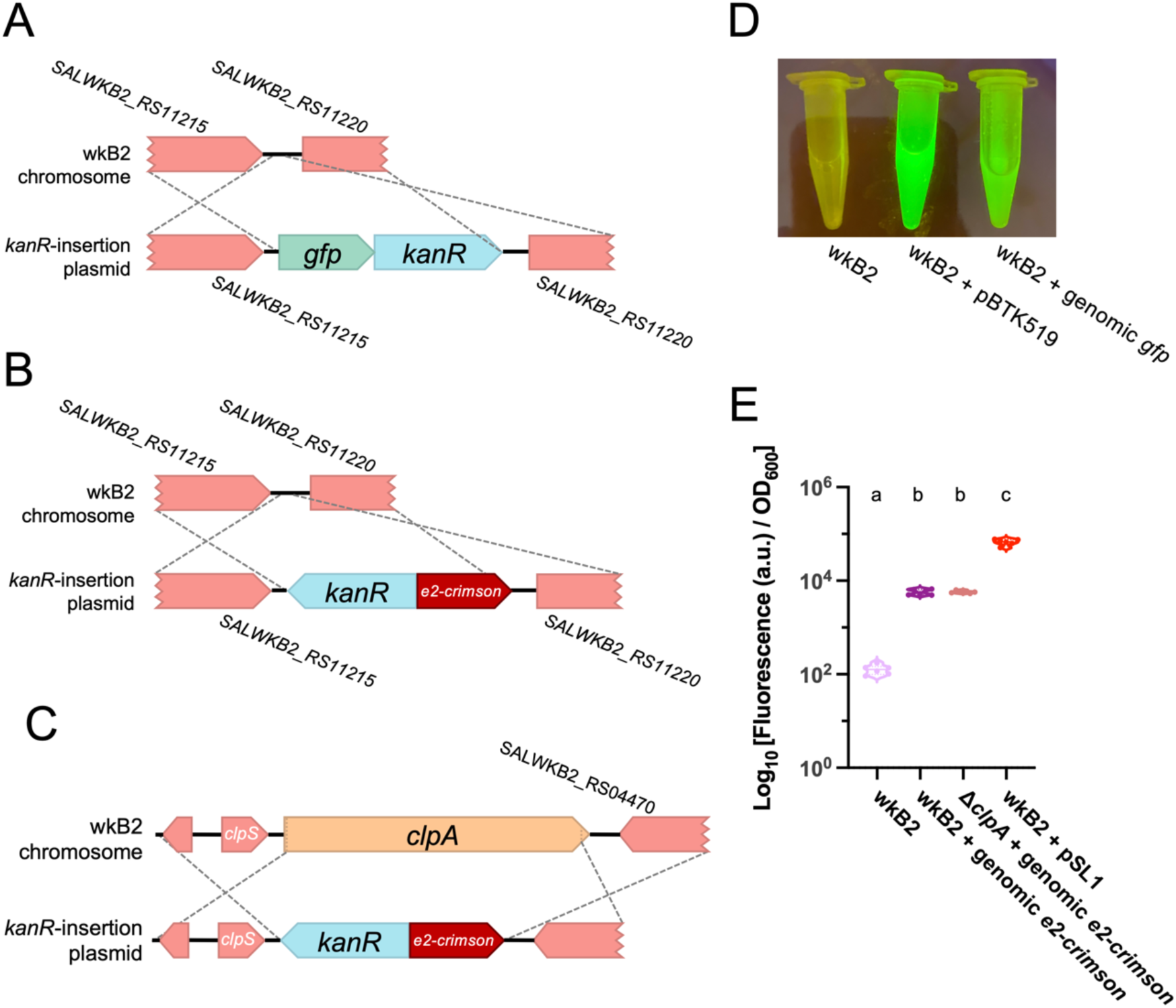
Fluorescent protein-coding genes can be inserted into the *S. alvi* genome. A. Gene insertion map depicting insertion of a *gfp* + *kanR* cassette into a non-essential, intergenic region of the genome. B. Gene insertion map depicting insertion of a *e2-crimson* + *kanR* cassette into a non-essential, intergenic region of the genome. C. Gene insertion map depicting deletion of the *clpA* gene via replacement insertion of a *e2-crimson* + *kanR* cassette. D. Image of wkB2, wkB2 + pBTK519 (positive control), and wkB2 + genomic *gfp* cells in front of a blue light transilluminator (470 nm), demonstrating successful insertion of *gfp* as evidenced by green fluorescence that is not observed in WT cells. E. Plot of quantification of *e2-crimson* fluorescence, demonstrating that wkB2 + genomic *e2-crimson* cell pellets resuspended in PBS display significantly increased red fluorescence compared to wkB2, and only slightly less fluorescence than wkB2 + pSL1, which contains a multicopy plasmid expressing *e2-crimson*. Log_10_ of OD_611 ex, 646 em_/OD_600_ is shown for cell cultures that were pelleted, resuspended in PBS, and read in a 96 well plate. Solid line = median; thin dotted line = quartiles. N=6 for each condition. Dissimilar letters above each group indicate a significant difference in means (P < 0.0001, as determined by One-way parametric ANOVA with Tukey’s multiple comparison test).

Next, we validated the phenotype of the *gfp* and *e2-crimson* genomic insertion strains. *S. alvi* wkB2 and wkB2 + genomic *gfp* cells were grown in liquid culture and imaged on a blue light transilluminator. wkB2 + genomic *gfp* were fluorescent, in contrast to wkB2 control cells (Fig 4D), indicating genomically-encoded *gfp* is capable of being successfully transcribed and translated into protein with observable fluorescence. To test expression of *e2-crimson*, liquid cultures of wkB2, wkB2 + genomic *e2-crimson*, and wkB2 + pSL1^36^ (which contains plasmid-encoded *e2-crimson*) were pelleted, resuspended in PBS, and quantified for E2-Crimson emission (622 nm excitation, 646 nm emission) normalized to cell growth (OD_600_). wkB2 + genomic *e2-crimson* had significantly higher fluorescence than wkB2 (Fig 4E), indicating successful production of genomically encoded E2-Crimson. Quantification of fluorescence in cultures prior to pelleting similarly indicated a significant increase in fluorescence in the mutant strains (Fig S8). As expected for one genomic copy versus a multicopy plasmid, wkB2 + genomic *e2-crimson* produced less fluorescence than did wkB2 + pSL1 (Fig 4E). These results indicate that exogenous genes can be successfully incorporated into the *S. alvi* genome.

### Gene knockout in *S. alvi* can be coupled to functional genomic insertion

Next we demonstrated that a gene knockout could be made at the same time as a functional genomic insertion. To this end, we combined insertion of a fluorescent protein-coding gene with deletion targeting the non-essential *clpA* gene. We designed a construct containing *e2-crimson* and *kanR*, flanked by arms with homology upstream and downstream of *clpA* (Fig 4C; Fig S7). Similar to cells containing a fluorescent protein-coding gene inserted into an intergenic region, we found that Δ*clpA* + *e2-crimson* cells emitted fluorescence, with no significant difference between these strains (Fig 4E). This result confirms that fluorescent protein-coding gene insertion can be combined with gene deletion.

### DNA integration is RecA-dependent

Having validated SnODIFY, we wanted to verify that genome integration was occurring through the expected RecA-mediated recombination mechanism that predominates in bacteria. To test this hypothesis, we generated a *recA::specR* strain and attempted to delete another gene, *amsE*, in this background using a *kanR* cassette. We found that, while *recA::specR* could still be transformed with a plasmid and *amsE* could be knocked out in wkB2, *amsE* could not similarly be knocked out in *recA::specR* cells (Fig. 5). To ensure that synthetic lethality between *asmE* and *recA* was not what accounted for lack of transformant, we repeated this experiment by trying to knock out *recJ* in a *recA::specR* background. As before, *recJ* could be knocked out in wild type *S. alvi* wkB2, but not in *recA::specR* cells (Fig. S9). This finding indicates that lack of double mutants in a *recA::specR* background is likely not due to synthetic lethality. Rather, these results demonstrate that gene knockout in *S. alvi* is RecA-dependent.

**Fig 5.**
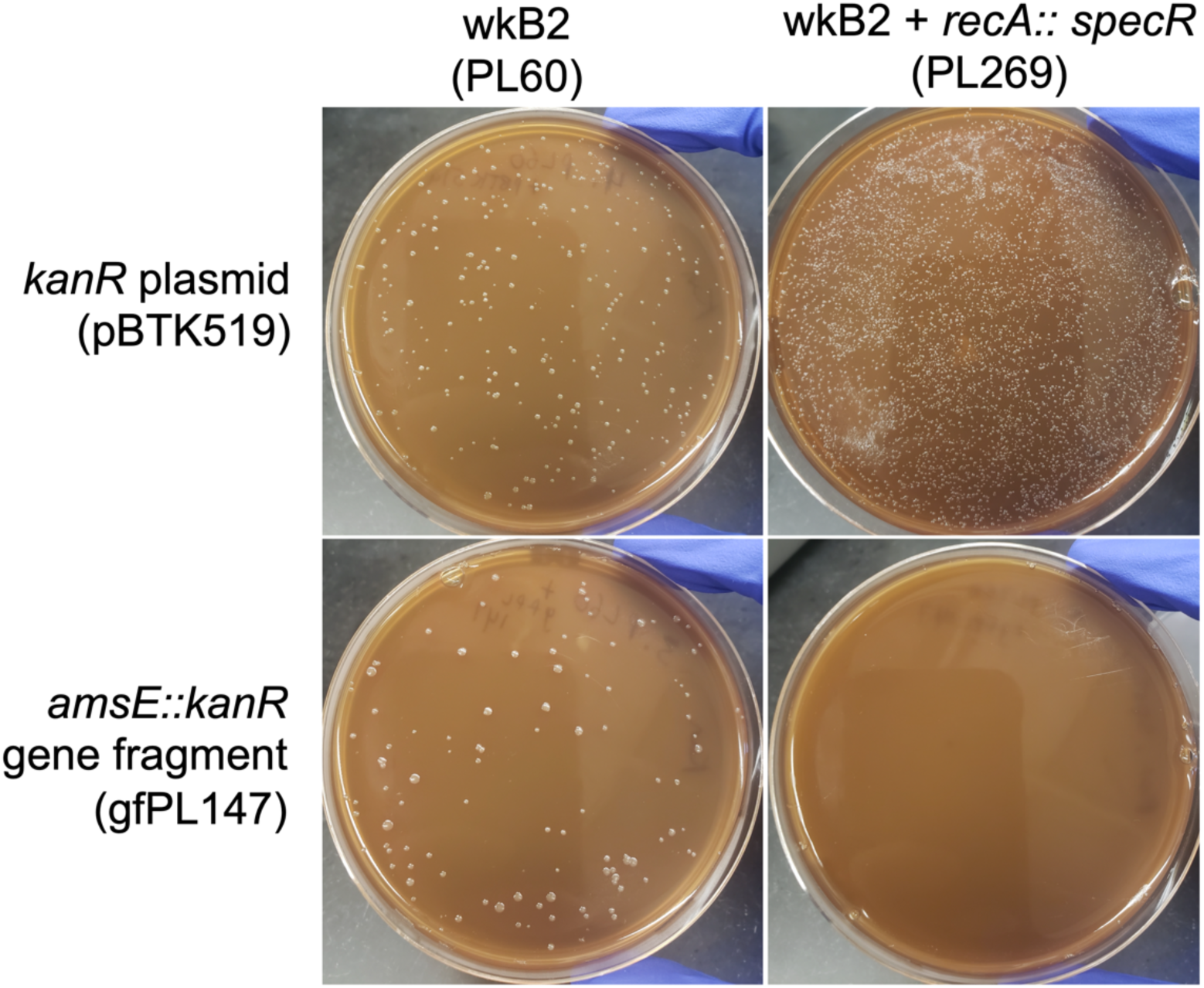
Gene knockout is *recA-*dependent. Plates demonstrating that wkB2 (PL60; top left) and wkB2 + *recA::specR* (PL269; top right) are successfully electroporated with a control *kanR* plasmid (pBTK519). The knockout construct *amsE::kanR* (gfPL147) is incorporated into wkB2 (bottom left), but not wkB2 + *recA::specR* (bottom right).

## Discussion

Here, we describe SnODIFY, a method for engineering the *S. alvi* chromosome using electroporation and homologous recombination. Strategies to genetically engineer other insect symbionts have had varied success^37^. While transposon mutagenesis (Tn5 and *Himar1*) has been described in a number of insect symbionts^21,37–40^, these strategies are not site-specific. Conversely, site-specific transposon mutagenesis via Tn7, while utilized in insect symbionts^37^, does not allow for gene deletion. Site-specific deletion has been demonstrated in a handful of insect symbionts, though these strategies have used expression of exogenous lambda Red machinery (BfO2, *S. glossinidius*)^41,42^, or use of a two-step suicide vector strategy (*R. rhodnii*, *S. glossinidius*, *B. insecticola*, *S. alvi*, *B. apis, F. perrara*)^22,37,38,41,43,44^, both of which add either complexity or time to knockout construction. Compared to symbiont engineering described in other insect systems, SnODIFY is highly accurate, straightforward, flexible, and reliable.

Integration of a circular plasmid into a genome is typically a two-step process^45^, with each crossover occurring sequentially. Previous modification of the *S. alvi* genome using conjugation of an R6K suicide vector resulted in mostly single-crossover integrations of the plasmid, and targeting Cas9 cleavage to the integration site was needed to increase the fraction of cells with double-crossover integration (i.e., just integration of the antibiotic resistance gene versus the entire plasmid)^22^. However, we observed only double-crossover integrants when we electroporated a ColE1 plasmid that cannot replicate in *S. alvi*, even when we did not linearize the plasmid. This discrepancy may be due to how conjugation delivers single-stranded DNA to a cell versus the dsDNA that was electroporated. Apparently, *S. alvi* recombination is efficient enough to favor the double-crossover integration, which makes linearization of the plasmid prior to electroporation unnecessary.

For each of the knockout strains we made, we observed what could be secondary mutations in the whole-genome sequencing results (Table S2; Table S3). While some of these mutations appear to already be present in our ancestral stock, others are found in only one or a few edited strains. Further work is needed to determine whether these mutations arose as suppressors to genes targeted for deletion, are common mutations in lab-cultured *S. alvi*, or were somehow selected for during the transformation/recombination process.

In its current form, SnODIFY provides immediate value to the bee microbiome field. This technique will facilitate study of *S. alvi*’s contribution to host-symbiont interactions by reverse genetics, allowing mechanistic investigation into host engraftment, immune stimulation, and host nutrient metabolism. SnODIFY will also enable investigations of how *S. alvi* interacts with other microbiome members and bee pathogens. Integration of fluorescent protein-coding genes into the genome of *S. alvi* allows it to be visualized in the bee host without needing to administer antibiotics to maintain a plasmid. Further, we foresee SnODIFY allowing for *S. alvi* engineering projects, ranging from migrating additional plasmid-based machinery to the genome, to construction of gain-of-function mutant strains that improve upon *S. alvi*’s probiotic capabilities^23,46^. Finally, the interoperability of SnODIFY and the bee microbiome toolkit (BTK)^22^ can expedite strain construction. Part plasmids constructed to contain homology arms can be mixed and matched with different payloads, allowing for streamlined knockout/insertion library construction.

With further development, new use cases and features could expand SnODIFY’s utility. Induction/depletion experiments (performed either with a gene knockout and complementation with a plasmid-based copy of a gene under control of an inducible promoter, or by placing an inducible promoter into the genome upstream of a gene) could be performed *in vivo*^47^, to study essential cellular processes on demand. Use of site-specific recombinases such as the Flp-FRT system for scarless gene deletion^48^ would remove confounders (such as the presence of antibiotic resistance cassettes) in attributing phenotypes to gene deletions and allow recycling antibiotic resistance cassettes to make multiple edits. Expression of exogenous recombineering systems (e.g., RecET or lambda Red) from a plasmid could improve recombination efficiency, as in *E. coli* or *Acinetobacter baumanii*, for example^35,49,50^. Finally, allelic exchange, facilitated by two-step recombination and use of counter-selectable markers^51^, would allow for study of loss-of-function or gain-of-function mutants. In short, we expect the SnODIFY platform presented in this paper to not only have immediate utility in reverse genetics and engineering applications in *S. alvi*, but to also serve as a springboard for future genome engineering toolsets.

## Methods

### Strains and DNA constructs

Strains, plasmids, oligos, and gene fragments are reported in Table S4.

### Bacterial cell culture

*S. alvi* cells were grown on Columbia + 5% sheep’s blood (Col-B) agar plates (with or without antibiotics, as indicated) at 35°C, 5% CO_2_, with 2-3 day incubation time. *S. alvi* cells were grown in liquid culture (with or without antibiotics, as indicated) at 35°C, 5% CO_2_, with 2-3 day incubation time. Media and containers used for liquid culture depended on the application, as indicated below. *E. coli* cells were grown with antibiotics on LB + 1.5% agar plates or in liquid LB, at 37°C, incubating overnight. Where indicated, 25 μg/mL kanamycin (kan) was used in *S. alvi* cultures; 30 μg/mL of spectinomycin (spec) was used in *S. alvi* cultures and 60 μg/mL of spectinomycin was used in *E. coli* cultures. Strains were saved in 30% glycerol stocks and stored at −80°C. Plasmids were obtained from *E. coli* overnight cultures using a Monarch plasmid miniprep kit (New England Biolabs, USA) and concentration was quantified by NanoDrop (Thermo Scientific, USA).

### Knockout construct design and synthesis

Putative genes identified for knockout were first determined to be non-essential *in vitro* by cross-referencing to the previously published *S. alvi* Tn-seq dataset^21^. Homology arms of 500 or 1000 bp either flanking or within the gene of interest were determined using NCBI gene, with wkB2 as a reference genome. Knockout constructs were then designed in Benchling’s molecular cloning tool^52^, by placing homology arms 5’ and 3’ to an antibiotic resistance cassette, either *kanR* or *specR*, flanked by a promoter and terminator. Constructs were commercially synthesized either as gBlocks (IDT, USA) or as plasmid inserts (Azenta, USA).

### Fluorescent protein-coding gene insertion construct design and synthesis

Insertion sites in the *S. alvi* genome were identified by finding two regions of the genome deemed non-essential by the *S. alvi* Tn-seq experiment^21^. The first was a 378 base-pair intergenic region between the genes *SALWKB2_RS11215* and *SALWKB2_RS11220*. The second was the non-essential gene *clpA*.

Golden Gate assembly part plasmids were made by cloning the parts into the cloning vector pBTK1001 using BsmBI-v2 (New England Biolabs, USA) (Fig S5; Fig S6). Flanking homology sequences of 1000 bp were PCR amplified from wkB2 genomic DNA with primers that added overhangs, so that they could be cloned into the entry vector pBTK1001 as Type 1 and Type 5 parts by BsmBI assembly, similar to previously described methods^22^. Integration cassettes were constructed as Type 2-3-4 parts in pBTK1001. Two versions of the Kanamycin resistance cassettes were amplified as Type 2 and Type 4 from pBTK1047. The E2-Crimson cassette was amplified as a Type 3-4 part from pLM70. The GFP cassette was amplified as Type 2-3 from pBTK503. Assembly reactions were heat-shock transformed into NEB 5-alpha *E. coli* cells (New England Biolabs, SUA) and plated on selective media. Transformants were cultured overnight. Part plasmids were extracted using a QIAprep Spin Miniprep Kit (Qiagen, Germany) and sequence verified by whole plasmid sequencing (Plasmidsaurus, USA). To create plasmids containing an integration cassette flanked by homology to the wkB2 genome, two homology parts and an integration cassette part were combined with the ColE1 origin-containing pYTK095 backbone^53^ using BsaI-HF v2 Golden Gate assembly (New England Biolabs, USA). As before, plasmids were transformed into *E. coli*, transformants selected for, and plasmid sequence verified.

### PCR amplification of knockout constructs

Synthesized DNA constructs were amplified via PCR. Briefly, primers with homology to the 5’ and 3’ ends of the construct were designed. For each construct, several 50 μl PCR reactions were run using these oligos and Q5 or Phusion polymerase (New England Biolabs, USA). A sample was run on a 1% agarose gel and imaged via UV transilluminator, to confirm successful amplification. Amplicons were purified using Magbeads (Axygen, USA) or a DNA Clean & Concentrator kit (Zymo Research, USA), and concentration was measured via NanoDrop (Thermo Scientific, USA). Constructs were brought to a suitable concentration to allow for electroporation with around 5 μg of DNA.

### *S. alvi* electroporation

Electrocompetent *S. alvi* cells were prepared prior to transformation in a manner similar to that commonly used for *E. coli*. Briefly, *S. alvi* was grown on solid media, as described above. After the appearance of colonies, cells were scraped and resuspended in a cell culture flask containing 50 ml Columbia broth. Cells were incubated without shaking at 35°C, 5% CO_2_ for 2-3 days. Working at room temperature, cells were then pelleted at approximately 5000 ×g for 10 minutes and supernatant decanted. Cells were washed twice by resuspending in 30 ml Ultrapure dH_2_O and pelleting, as above. After the third centrifugation, cells were resuspended in 1 ml filtered 10% glycerol in dH_2_O, dispensed into 100 μl aliquots, and stored at −80°C.

*S. alvi* cells were electroporated in a manner similar to that used for *E. coli*, with a few modifications. Briefly, electrocompetent cell preps were thawed on ice and incubated with approximately 500 ng of plasmid DNA or 5-10 μg of knockout/insertion construct DNA for 20 minutes (5 μg of DNA was used to construct most knockout strains; 10 μg of DNA was used to construct PL269 and PL271). The cell + construct mixture was added to either a 1 mm or 2 mm electroporation cuvette depending on the volume. Cells were electroporated with a Gene Pulser Xcell Electroporator (Bio-Rad, USA) using 1.8 kV (for 1 mm cuvettes) or 3 kV (for 2 mm cuvettes). Cells were resuspended in Columbia broth and allowed to recover without antibiotic overnight at 35°C, 5% CO_2_. The next day, cells were pelleted, resuspended in 100 μl of media, and the entirety plated on Columbia + 5% sheep’s blood agar plates with the respective antibiotic. Plates were incubated for 2-3 days at 35°C, 5% CO_2,_ until the appearance of colonies. Multiple colonies were then isolated by passage onto a fresh Columbia + 5% sheep’s blood agar plate, which were allowed to form colonies.

### PCR screen for knockout mutants

Colony PCR was performed to screen for proper genomic insertion of the knockout/insertion construct. Briefly, 2-14 colonies were picked from the transformation isolation plates and resuspended in 20 μl of dH_2_O. PCR was run using diluted template, oligos flanking the antibiotic resistance cassette, and Q5 polymerase. A control PCR was also run using WT wkB2. A portion of the PCR reaction was run on a 1% agarose gel and imaged using a UV transilluminator to visualize DNA band size. Amplicons of interest were prepared using magnetic beads (Axygen, USA) and concentration was quantified via NanoDrop (Thermo Scientific, USA). Amplicons were sent for Sanger or Next-gen amplicon sequencing. Sequences were aligned to the designed knockout construct reference using Benchling’s molecular cloning tool to confirm presence of the antibiotic resistance cassette.

### Knockout/insertion confirmation by WGS

We sequenced strains with genome modifications using either short-read (Illumina) or long-read (Nanopore) technologies. For short-read sequencing, genomic DNA (gDNA) was purified from the relevant strain using a DNeasy Blood & Tissue kit (Qiagen, Germany). For long-read sequencing, high molecular weight gDNA was isolated using the Quick-DNA HMW MagBead kit (Zymo Research, USA). gDNA concentrations were measured by Qubit (Invitrogen, USA) or NanoDrop (Thermo Scientific, USA).

gDNA was then sequenced in one of three ways: 1. Commercial library preparation and short-read sequencing (SeqCenter, USA); 2. In-house long-read sequencing; or 3. In-house library preparation and commercial short-read sequencing (Psomagen, USA). Commercial library preparation used the tagmentation DNA Prep Kit (Illumina, USA). For in-house short-read sequencing library preparation, 10 ng gDNA was inputted into the xGen DNA lib Prep EZ kit using the xGen Deceleration module (Integrated DNA Technologies, USA). All reactions were carried out at 20% of the manufacturer’s recommended volumes, with dual 6-bp indexes incorporated during a final 12 cycle PCR step. For long-read sequencing, 175 ng of gDNA was prepared using a Rapid Barcoding Sequencing Kit and sequenced on a R9.4.1 flow cell using a MinION MK1C instrument (Oxford Nanopore, UK). Basecalling was done using Guppy (v6.3.9) in fast mode. Raw FASTQ files are available from the NCBI Sequence Read Archive (PRJNA1017617).

Long-read FASTQ files were trimmed using Porechop (v0.2.4)^54^ with the option to discard reads with internal adaptors. Short-read FASTQ files were trimmed using fastp (v.0.23.4)^55^. Variant calling was performed on trimmed FASTQ files using *breseq* (v0.38.1)^56^ with the *S. alvi* wkB2 genome (Genbank: CP007446)^33^ and the insertion deletion/cassette sequences as references. Mutations, including the deletion/resistance cassette insertion and secondary mutations, were annotated. For analysis of Nanopore data, indels in homopolymers of 3+ bases were not reported.

### Crystal violet biofilm formation assay

Biofilm formation in *S. alvi* was assayed using a Crystal violet assay, as previously described^21,35^. Briefly, *S. alvi* strains were inoculated directly from a freezer stock into an overnight culture in a 50 ml conical containing Insectagro DS2 (Corning, USA) + antibiotic, and incubated for 3 days at 35°C, 5% CO_2_. 1:40 dilutions of confluent cells (including resuspended biofilm) were made in Insectagro DS2 without antibiotics in a 96 well plate. (Note that outer wells were filled with either blank media or culture that would not be used in downstream analysis, due to differential growth resulting from differential oxygen availability). Plates were incubated for 2 days at 35°C, 5% CO_2_. After 2 days, OD_600_ was measured on a Spark 10M plate reader (Tecan, Switzerland) to quantify overall cell growth. Plates were washed 3 times by dunking in dH_2_O and stained in 0.1 % Crystal violet (in dH_2_O) for 10 minutes. Stain was then removed and plates were allowed to dry in a warm room (37°C) for 1-3 hours. Dried plates were then imaged. 30% acetic acid (in dH_2_O) was added to resolubilize the dried Crystal violet and incubated for 10 minutes. OD_550_ was then measured on a Spark 10M to quantify the amount of biofilm present.

Plate reading data were analyzed by first subtracting background (blank reading) in the OD_600_ and OD_550_ readings. Outer wells were omitted from analysis (see above) and OD_600_ values of less than 0.01 were removed. OD_550_ was then divided by OD_600_, and OD_550_/OD_600_ was plotted in Prism Graphpad for each strain. Only positive OD_550_/OD_600_ are displayed. A Shapiro-Wilk Normality test was run to determine if each sample was normally distributed and each condition passed. A One-way parametric ANOVA with Dunnett’s multiple comparison test (comparing to the WT control) was subsequently performed, and significance was indicated on plots.

### Imaging and quantification of fluorescent strains

Fluorescent insertion strains were imaged and/or quantified for fluorescence. wkB2 + genomic *gfp* and wkB2 were grown for 2 days on Columbia agar + 5% Sheep’s blood + kanamycin media at 35 °C 5% CO_2_. After 2 days, cultures were scraped, resuspended in PBS in microcentrifuge tubes, and imaged in front of a blue light transilluminator (470 nm). For experiments involving E2-Crimson, wkB2 + genomic *e2-crimson*, wkB2 + pSL1, and wkB2 were grown for 2 days in liquid BHI media in a sealed bag maintaining a microaerobic environment (85% nitrogen; 10% CO_2_; 5% O_2_)^57^. After 2 days, cultures were pelleted and imaged. Pellets were then resuspended in PBS and transferred to a 96 well plate. Samples were read in a Spark 10M plate reader in triplicate, using 611 nm excitation and 646 nm emission^58^ and optimal gain was determined by the plate reader. Absorbance readings from technical replicates were averaged, then log-transformed and plotted in GraphPad Prism (Dotmatics, USA), with individual data points for each group representing biological replicates. A Shapiro-Wilk Normality test was run to determine if each sample was normally distributed, and each condition passed. A one-way parametric ANOVA with Tukey’s multiple comparison test was subsequently performed and significance indicated on plots.

## Acknowledgements

We thank Daniel Deatherage for performing whole-genome sequencing. We also thank Eli Powell for technical advice, as well as members of the Moran and Barrick lab for discussions. Figures were made using BioRender. This work was supported by funding from the National Science Foundation (IOS-2103208), U.S. Army Research Office (W911NF-20-1-0195), National Institutes of Health (R35GM131738), and USDA NIFA (2023-67012-39356).

## Author contributions

Conceptualization, P.J.L., S.P.L., A.H.M.Z.A., N.A.M., and J.E.B.; Methodology, P.J.L., S.P.L., and A.H.M.Z.A.; Investigation, P.J.L., A.H.M.Z.A., and L.G.M.; Writing, P.J.L. and A.H.M.Z.A.; Editing, P.J.L., S.P.L., A.H.M.Z.A., N.A.M., and J.E.B.

## Supplemental tables and figures

**Table S1-S4: See supplemental attachments**

**Fig S1.**
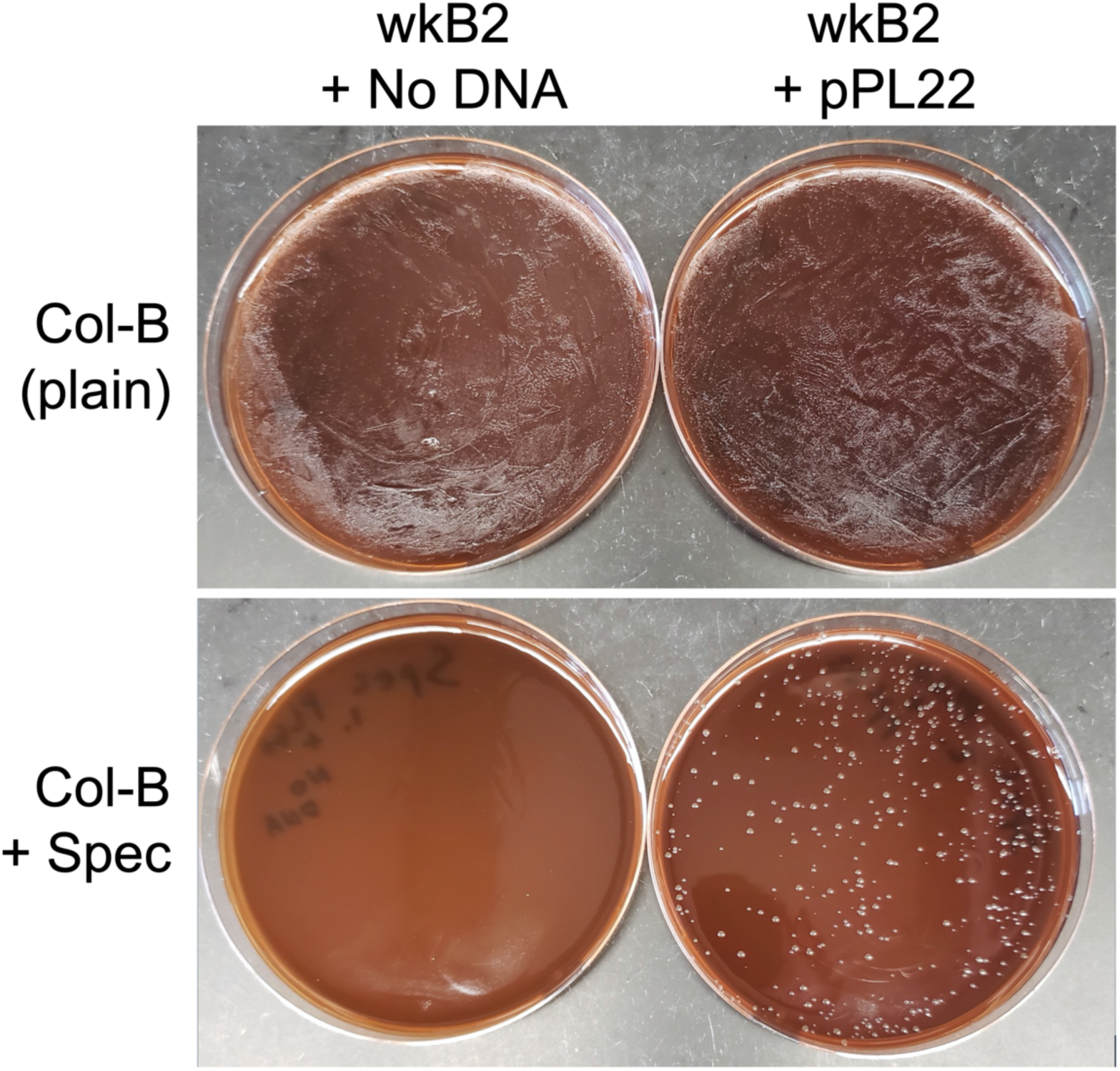
*S. alvi* can be transformed by electroporation. Images of transformation plates demonstrating *S. alvi* can be transformed with a plasmid by electroporation. *S. alvi* that has received a *specR*-containing plasmid (wkB2 + pPL22, bottom right) is able to grow on Col-B agar + Spec plates, whereas wkB2 + no DNA (bottom left) cannot. wkB2 transformation reactions with pPL22 (top right) or no DNA (top left) plated on plain Col-B demonstrate that transformation reactions yield viable cells.

**Fig S2.**
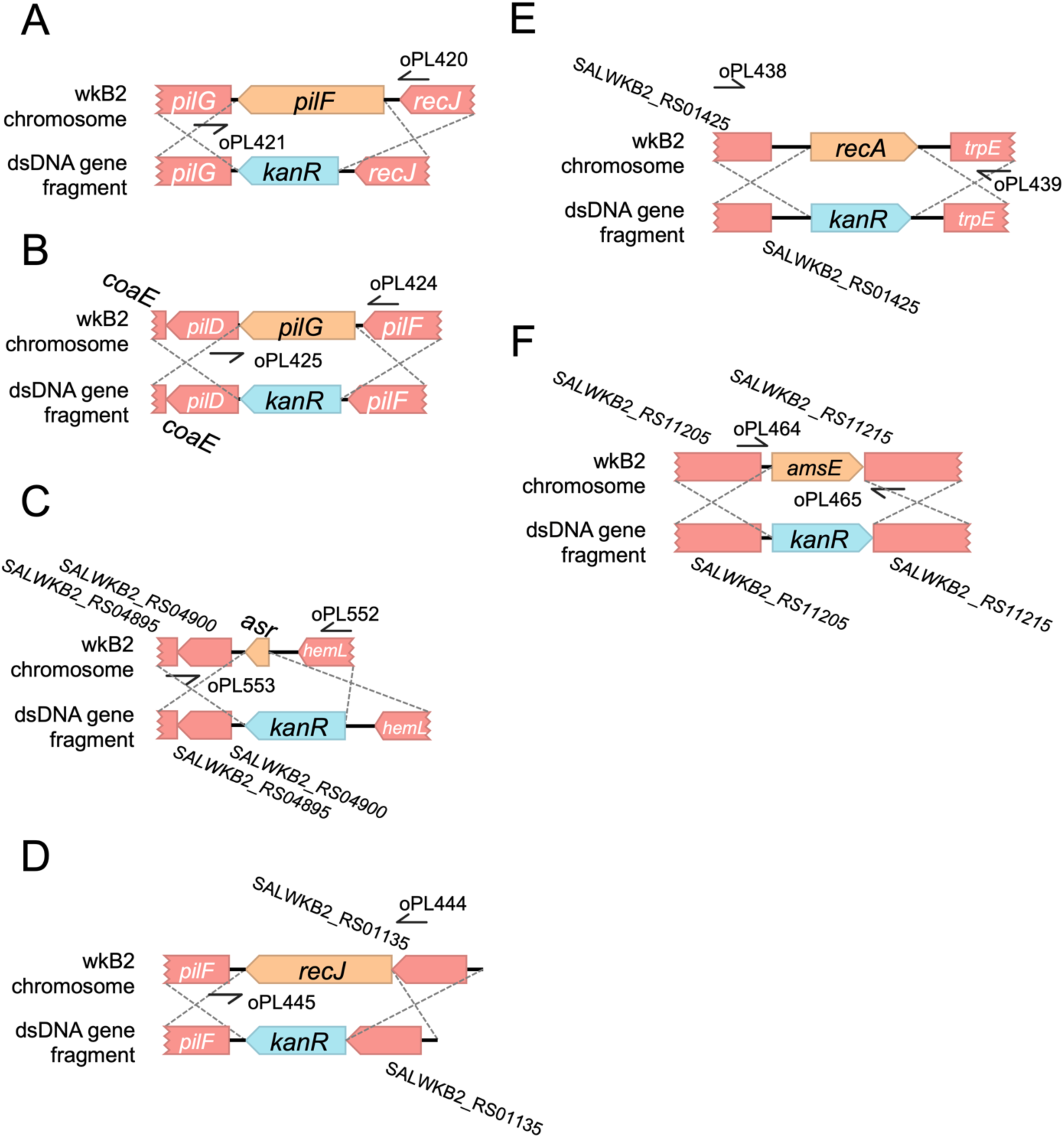
Construct design of additional gene knockouts. A,B,C,D,E,F. Deletion maps of indicated genes in their genomic contexts: *pilF* (A), *pilG* (B), *asr* (C), *recJ* (D), *recA* (E), *amsE* (F). For each gene, deletion construct and screening primers are depicted.

**Fig S3.**
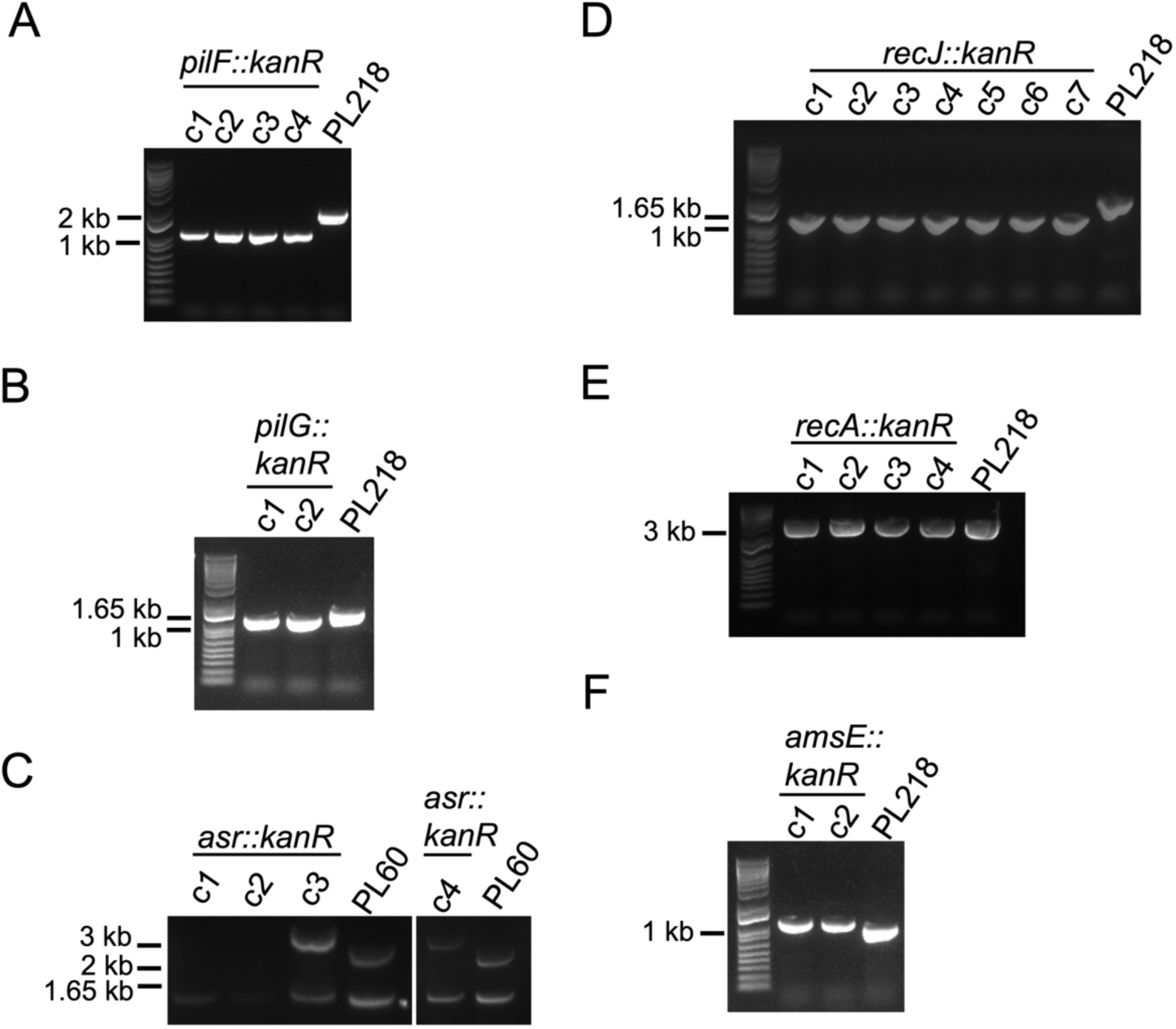
Knockout of nearly all additional genes is highly accurate. DNA gels of PCR screen of additional knockout strains. PCR products of clones of each knockout and a WT control [PL60 (wkB2) or PL218 (wkB2 + pDS-GFP)] were run with the corresponding primers (shown in Fig S2.) A. A 1000 bp band corresponding with *kanR* is observed in 4 out of 4 *pilF::kanR* clones, compared to the 2000 bp control band, indicating 100% editing accuracy. B. A 1000 bp band corresponding with *kanR* is observed in 2 out of 2 *pilG::kanR* clones, compared to the 2000 bp control band, indicating 100% editing accuracy. C. A ∼3000 bp band corresponding with *kanR* + flanking regions is observed in 2 of the 4 tested *pilG::kanR* clones, compared to the 2207 bp control, indicating 50% editing accuracy. Note, a ∼1500 bp off-target band is seen in both experimental and control sample PCR reactions, likely arising due to inefficient PCR. D. A 1000 bp band corresponding with *kanR* is observed in 7 out of 7 *recJ::kanR* clones, compared to the 1755 bp control band, indicating 100% editing accuracy. E. A ∼3000 bp band is observed in *recA::kanR* clones and control sample. As both *kanR* and control bands are predicted to be ∼3000 bp, editing accuracy is not evident from PCR alone. F. A band just over 1000 bp corresponding with *kanR* is observed in 2 out of 2 *amsE::kanR* clones, compared to the 922 bp control band, indicating 100% editing accuracy.

**Fig S4.**
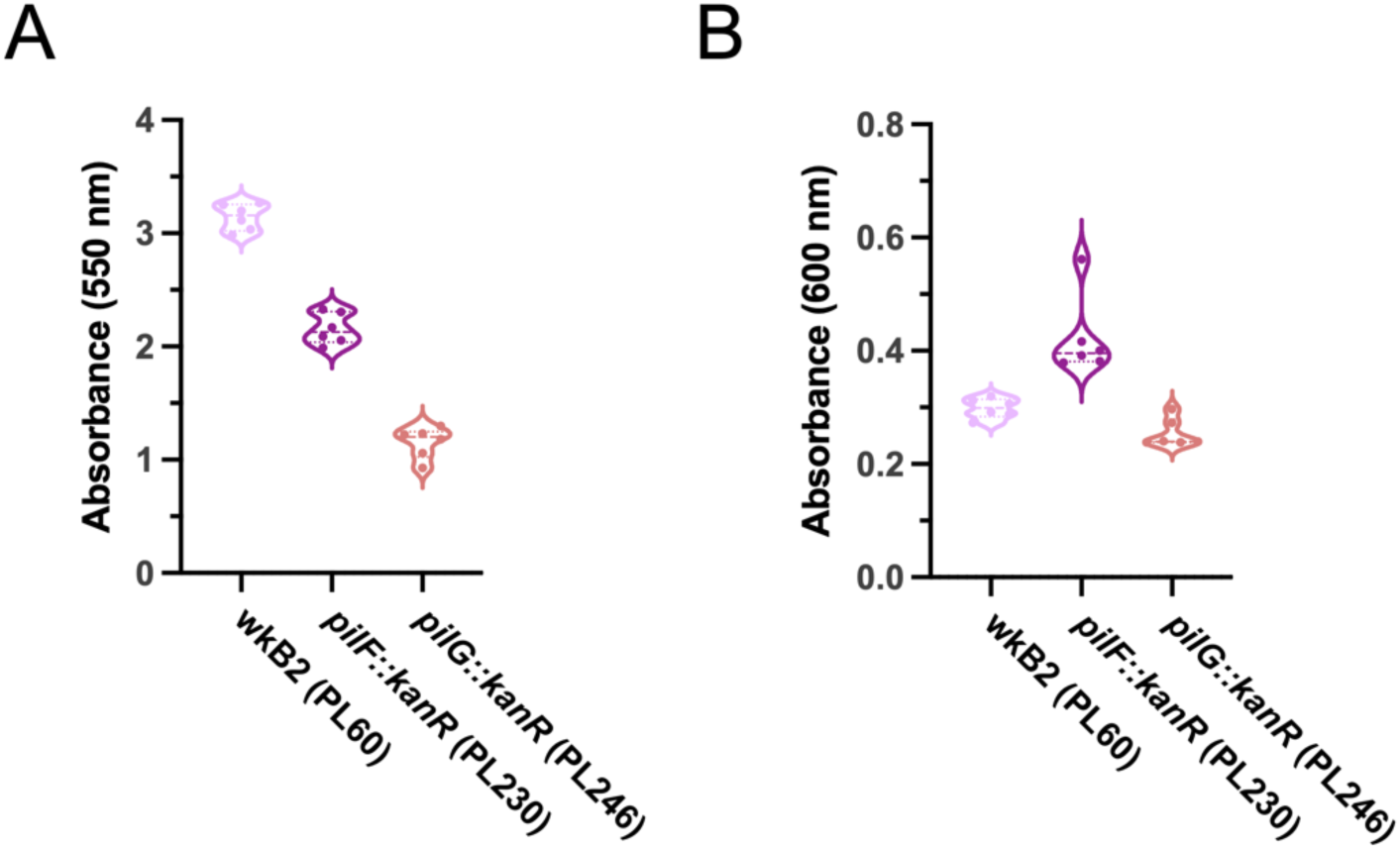
Biofilm formation is impaired in mutants, with limited impact to growth. A. Plot of OD_550_ to quantify biofilm formation for wkB2 and biofilm knockout strains, demonstrating that *pilF::kanR* and *pilG::kanR* have decreased biofilm formation. B. Plot of OD_600_ to quantify cell growth for wkB2 and biofilm knockout strains, demonstrating minor differences in growth. For A and B, OD_550_ and OD_600_ values were used to calculate OD_550_/OD_600_ in Fig 3A. Thick dotted line = median; thin dotted line = quartiles. N=6 for each condition.

**Fig S5.**
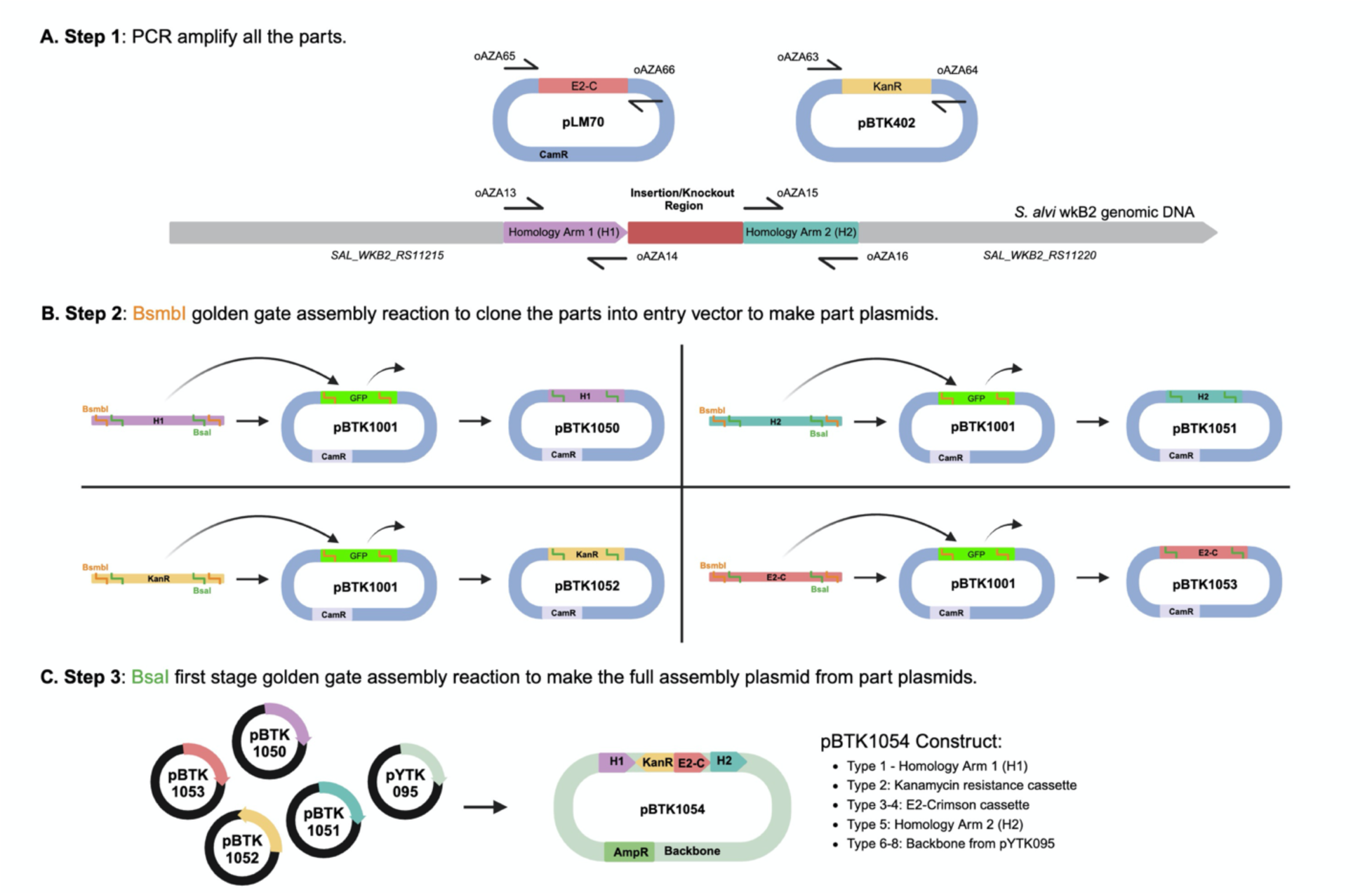
*e2-crimson* genomic insertion plasmid assembly diagram. A. Plasmid and genome maps depicting regions that were used as templates for insert construction. *e2-crimson* (from pLM70), *kanR* (from pBTK402), and homology arms flanking the insertion site (from *S. alvi* genomic DNA) were amplified via PCR. Primers used to amplify these parts contain BsmbI and BsaI recognition and cut sites. B. Plasmid maps depicting assembly of part plasmids. Parts were cloned into entry vectors (pBTK1001) using BsmbI Golden Gate Assembly mix (NEB, USA) to make part plasmids containing homology arms, *e2-crimson*, and *kanR*. C. Plasmid maps depicting final assembly of the part plasmids (Type 1-8) using BsaI first-stage Golden Gate Assembly mix (NEB, USA) to create the *e2-crimson* genomic insertion plasmid.

**Fig S6.**
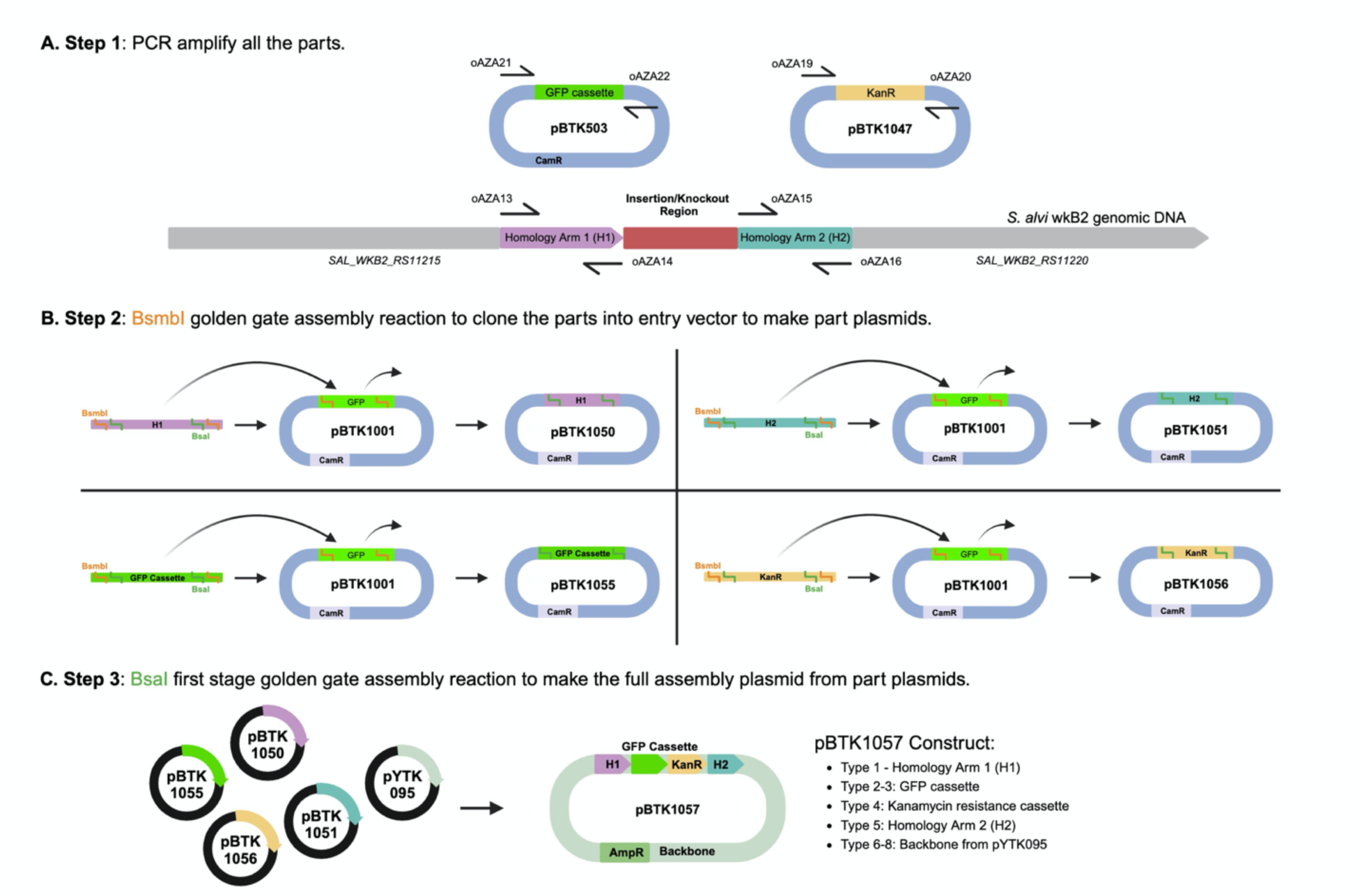
*gfp* genomic insertion plasmid assembly diagram. A. Plasmid and genome maps depicting regions that were used as templates for insert construction. *gfp* (from pBTK503), *kanR* (from pBTK1047), and homology arms flanking the insertion site (from *S. alvi* genomic DNA) were amplified via PCR. Primers used to amplify these parts contain BsmbI and BsaI recognition and cut sites. B. Plasmid maps depicting assembly of part plasmids. Parts were cloned into entry vectors (pBTK1001) using BsmbI Golden Gate Assembly mix (NEB, USA) to make part plasmids containing homology arms, *gfp*, and *kanR*. C. Plasmid maps depicting final assembly of the part plasmids (Type 1-8) using BsaI first-stage Golden Gate Assembly mix (NEB, USA) to create the *gfp* genomic insertion plasmid.

**Fig S7.**
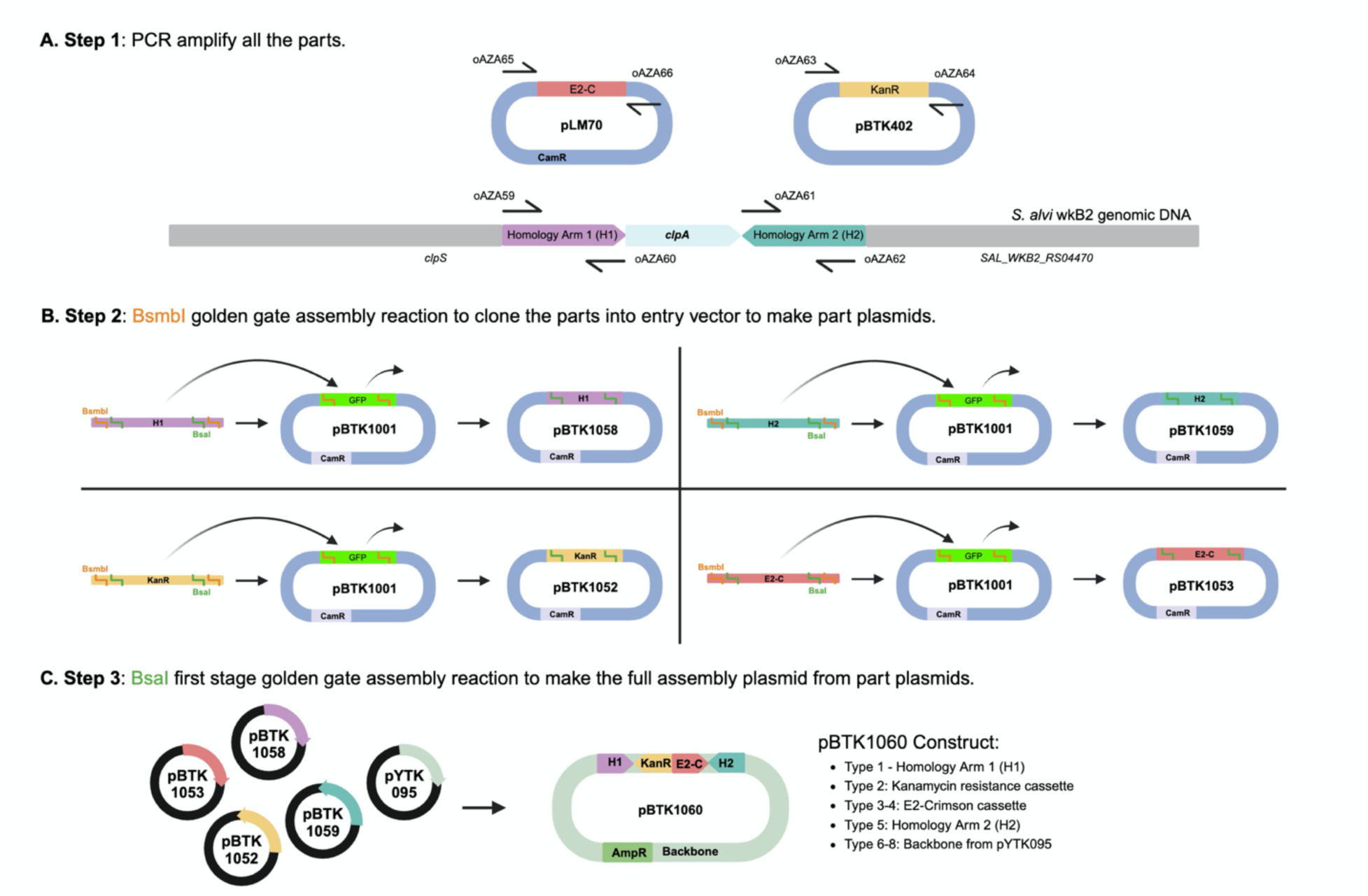
Δ*clpA* + *e2-crimson* genomic insertion plasmid assembly diagram. A. Plasmid and genome maps depicting regions that were used as templates for insert construction. *e2-crimson* (from pML70), *kanR* (from pBTK402), and homology arms flanking *clpA* (from *S. alvi* genomic DNA) were amplified via PCR. Primers used to amplify these parts contain BsmbI and BsaI recognition and cut sites. B. Plasmid maps depicting assembly of part plasmids. Parts were cloned into entry vectors (pBTK1001) using BsmbI Golden Gate Assembly mix (NEB, USA) to make part plasmids containing homology arms, *e2-crimson*, and *kanR*. C. Plasmid maps depicting final assembly of the part plasmids (Type 1-8) using BsaI first-stage Golden Gate Assembly mix (NEB, USA) to create the Δ*clpA* + *e2-crimson* genomic insertion plasmid.

**Fig S8.**
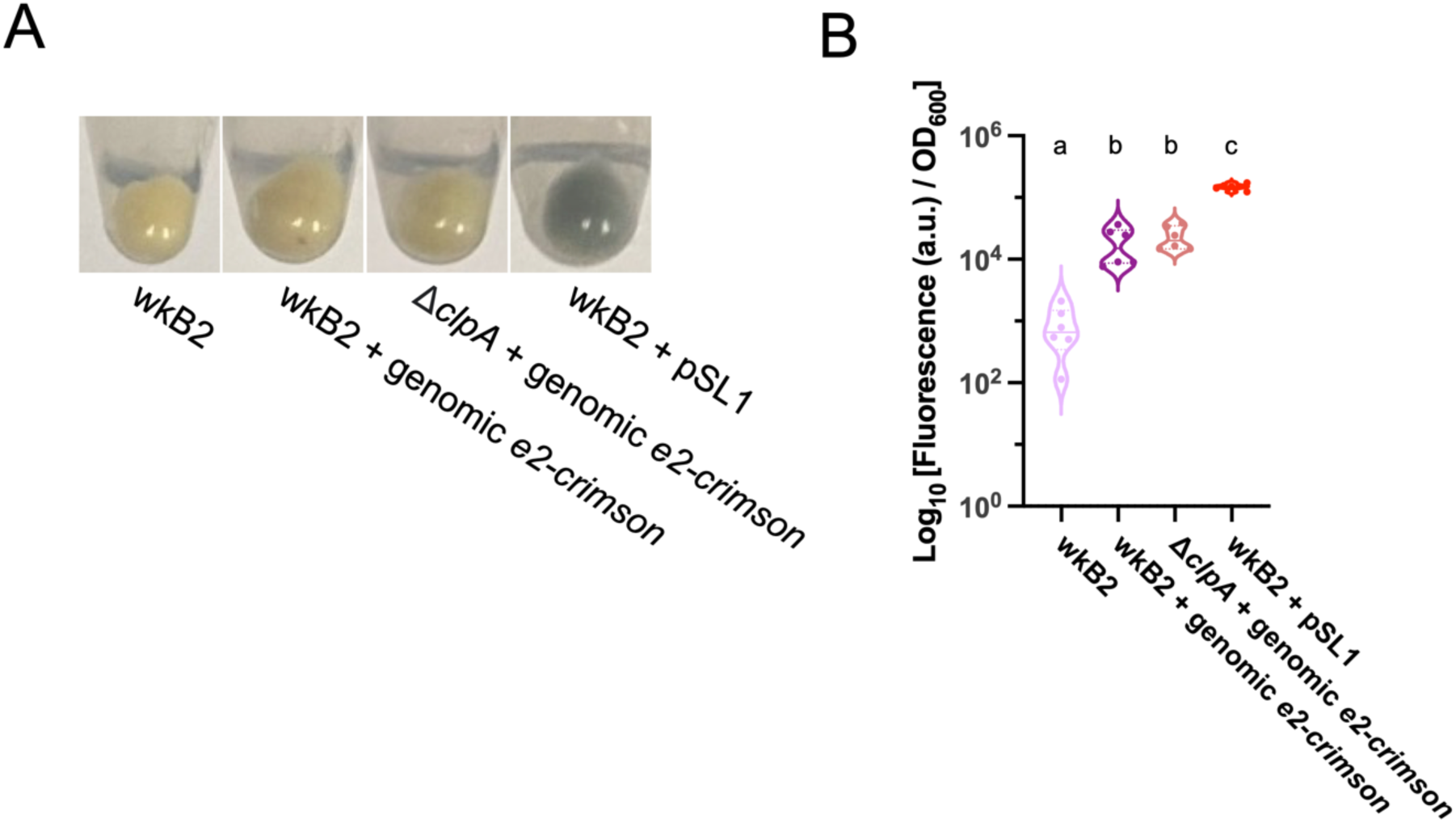
Genomic insertion of *e2-crimson* leads to increase in culture fluorescence. A. Image of pelleted wkB2, wkB2 + pSL1, wkB2 + genomic *e2-crimson*, and Δ*clpA* + *e2-crimson* cultures, demonstrating successful genomic insertion of *e2-crimson* induces a slight increase in purple hue, compared to the darker purple wkB2 + pSL1 and pale wkB2. B. Plot of quantification of *e2-crimson* fluorescence, demonstrating that wkB2 + genomic *e2-crimson* cultures grown in BHI display significantly increased red fluorescence compared to wkB2, and only slightly less fluorescence than wkB2 + pSL1, which contains multiple plasmid copies of *e2-crimson*. Log_10_ of OD_611 ex, 646 em_/OD_600_ is shown for cell cultures grown in BHI that were read in a 96 well plate. Solid line = median; thin dotted line = quartiles. N=6 for each condition. Dissimilar letters above each group indicate a significant difference in means (P < 0.0005, as determined by One-way parametric ANOVA with Tukey’s multiple comparison test).

**Fig S9.**
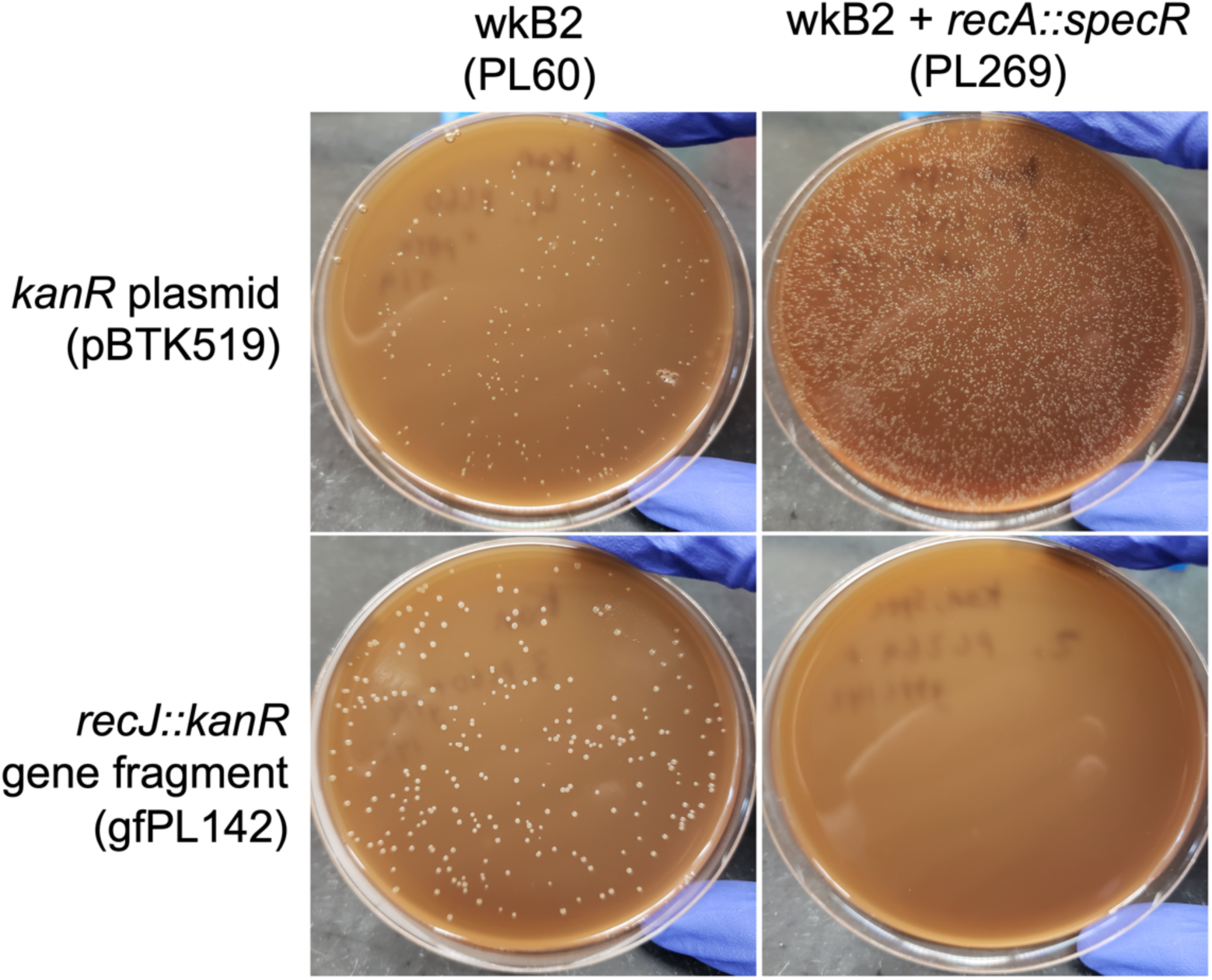
Knockout of additional gene, *recJ*, is not possible in a *recA::specR* background. Plates demonstrating that wkB2 (PL60; top left) and wkB2 + *recA::specR* (PL269; top right) are successfully electroporated with a control *kanR* plasmid (pBTK519). The knockout construct *recJ::kanR* (gfPL142) is incorporated into wkB2 (bottom left), but not wkB2 + *recA::specR* (bottom right).

## References

1. Bailes, E. J., Ollerton, J., Pattrick, J. G. & Glover, B. J. How can an understanding of plant– pollinator interactions contribute to global food security? Curr. Opin. Plant Biol. 26, 72–79 (2015).

2. Marshman, J., Blay-Palmer, A. & Landman, K. Anthropocene Crisis: Climate Change, Pollinators, and Food Security. Environments 6, 22 (2019).

3. van der Sluijs, J. P. & Vaage, N. S. Pollinators and Global Food Security: the Need for Holistic Global Stewardship. Food Ethics 1, 75–91 (2016).

4. Genersch, E. American Foulbrood in honeybees and its causative agent, *Paenibacillus larvae*. J. Invertebr. Pathol. 103, S10–S19 (2010).

5. Genersch, E. Honey bee pathology: current threats to honey bees and beekeeping. Appl. Microbiol. Biotechnol. 87, 87–97 (2010).

6. Report on the national stakeholders conference on honey bee health. 9–68 (2013).

7. Kwong, W. K. & Moran, N. A. Gut microbial communities of social bees. Nat. Rev. Microbiol. 14, 374–384 (2016).

8. Raymann, K. & Moran, N. A. The role of the gut microbiome in health and disease of adult honey bee workers. Curr. Opin. Insect Sci. 26, 97–104 (2018).

9. Zheng, H., Powell, J. E., Steele, M. I., Dietrich, C. & Moran, N. A. Honeybee gut microbiota promotes host weight gain via bacterial metabolism and hormonal signaling. Proc. Natl. Acad. Sci. U. S. A. 114, 4775–4780 (2017).

10. Raymann, K., Shaffer, Z. & Moran, N. A. Antibiotic exposure perturbs the gut microbiota and elevates mortality in honeybees. PLoS Biol. 15, e2001861 (2017).

11. Kešnerová, L. et al. Disentangling metabolic functions of bacteria in the honey bee gut. PLoS Biol. 15, e2003467 (2017).

12. Bonilla-Rosso, G. & Engel, P. Functional roles and metabolic niches in the honey bee gut microbiota. Curr. Opin. Microbiol. 43, 69–76 (2018).

13. Martinson, V. G., Moy, J. & Moran, N. A. Establishment of characteristic gut bacteria during development of the honeybee worker. Appl. Environ. Microbiol. 78, 2830–2840 (2012).

14. Steele, M. I., Motta, E. V. S., Gattu, T., Martinez, D. & Moran, N. A. The Gut Microbiota Protects Bees from Invasion by a Bacterial Pathogen. Microbiol. Spectr. 9, e0039421 (2021).

15. Kwong, W. K., Mancenido, A. L. & Moran, N. A. Immune system stimulation by the native gut microbiota of honey bees. R. Soc. Open Sci. 4, 170003 (2017).

16. Huang, Q., Lariviere, P. J., Powell, J. E. & Moran, N. A. Engineered gut symbiont inhibits microsporidian parasite and improves honey bee survival. Proc. Natl. Acad. Sci. 120, e2220922120 (2023).

17. Leonard, S. P. et al. Engineered symbionts activate honey bee immunity and limit pathogens. Science 367, 573–576 (2020).

18. Lariviere, P. J., Leonard, S. P., Horak, R. D., Powell, J. E. & Barrick, J. E. Honey bee functional genomics using symbiont-mediated RNAi. Nat. Protoc. 18, 902–928 (2023).

19. Lang, H. et al. Engineered symbiotic bacteria interfering *Nosema* redox system inhibit microsporidia parasitism in honeybees. Nat. Commun. 14, 2778 (2023).

20. Horak, R. D., Leonard, S. P. & Moran, N. A. Symbionts shape host innate immunity in honeybees. Proc. Biol. Sci. 287, 20201184 (2020).

21. Powell, J. E., Leonard, S. P., Kwong, W. K., Engel, P. & Moran, N. A. Genome-wide screen identifies host colonization determinants in a bacterial gut symbiont. Proc. Natl. Acad. Sci. 113, 13887–13892 (2016).

22. Leonard, S. P. et al. Genetic engineering of bee gut microbiome bacteria with a toolkit for modular assembly of broad-host-range plasmids. ACS Synth. Biol. 7, 1279–1290 (2018).

23. Powell, J. E., Carver, Z., Leonard, S. P. & Moran, N. A. Field-Realistic Tylosin Exposure Impacts Honey Bee Microbiota and Pathogen Susceptibility, Which Is Ameliorated by Native Gut Probiotics. Microbiol. Spectr. 9, e0010321 (2021).

24. Fels, U., Gevaert, K. & Van Damme, P. Bacterial Genetic Engineering by Means of Recombineering for Reverse Genetics. Front. Microbiol. 11, (2020).

25. Kurushima, J. & Tomita, H. Advances of genetic engineering in streptococci and enterococci. Microbiol. Immunol. 66, 411–417 (2022).

26. Kormanec, J. et al. Recent achievements in the generation of stable genome alterations/mutations in species of the genus *Streptomyces*. Appl. Microbiol. Biotechnol. 103, 5463–5482 (2019).

27. Sharan, S. K., Thomason, L. C., Kuznetsov, S. G. & Court, D. L. Recombineering: a homologous recombination-based method of genetic engineering. Nat. Protoc. 4, 206–223 (2009).

28. Court, D. L., Sawitzke, J. A. & Thomason, L. C. Genetic engineering using homologous recombination. Annu. Rev. Genet. 36, 361–388 (2002).

29. Kwong, W. K. & Moran, N. A. Cultivation and characterization of the gut symbionts of honey bees and bumble bees: Description of *Snodgrassella alvi* gen. nov., sp. nov., a member of the family *Neisseriaceae* of the *Betaproteobacteria*, and *Gilliamella apicola* gen. nov., sp. nov., a member of *Orbaceae* fam. nov., *Orbales* ord. nov., a sister taxon to the order ‘*Enterobacteriales*’ of the *Gammaproteobacteria*. Int. J. Syst. Evol. Microbiol. 63, 2008–2018 (2013).

30. van Dam, V. & Bos, M. P. Generating knock-out and complementation strains of *Neisseria meningitidis*. in Neisseria meningitidis: Advanced Methods and Protocols (ed. Christodoulides, M.) 55–72 (Humana Press, 2012). doi:10.1007/978-1-61779-346-2_4.

31. Aranda, J., et al. *Acinetobacter baumannii* RecA protein in repair of DNA damage, antimicrobial resistance, general stress response, and virulence. J. Bacteriol. 193, 3740–3747 (2011).

32. Koomey, M., Gotschlich, E. C., Robbins, K., Bergström, S. & Swanson, J. Effects of *recA* mutations on pilus antigenic variation and phase transitions in *Neisseria gonorrhoeae*. Genetics 117, 391–398 (1987).

33. Kwong, W. K., Engel, P., Koch, H. & Moran, N. A. Genomics and host specialization of honey bee and bumble bee gut symbionts. Proc. Natl. Acad. Sci. 111, 11509–11514 (2014).

34. Gonzales, M. F., Brooks, T., Pukatzki, S. U. & Provenzano, D. Rapid protocol for preparation of electrocompetent *Escherichia coli* and *Vibrio cholerae*. J. Vis. Exp. JoVE 50684 (2013) doi:10.3791/50684.

35. Tucker, A. T. et al. Defining Gene-Phenotype Relationships in *Acinetobacter baumannii* through One-Step Chromosomal Gene Inactivation. mBio 5, e01313–14 (2014).

36. Li, Y., Leonard, S. P., Powell, J. E. & Moran, N. A. Species divergence in gut-restricted bacteria of social bees. Proc. Natl. Acad. Sci. 119, e2115013119 (2022).

37. Elston, K. M. et al. Engineering insects from the endosymbiont out. Trends Microbiol. 30, 79–96 (2022).

38. Kendra, C. G., Keller, C. M., Bruna, R. E. & Pontes, M. H. Conjugal DNA Transfer in *Sodalis glossinidius*, a Maternally Inherited Symbiont of Tsetse Flies. mSphere 5, 10.1128/msphere.00864-20 (2020).

39. Wang, S. et al. Driving mosquito refractoriness to *Plasmodium falciparum* with engineered symbiotic bacteria. Science 357, 1399–1402 (2017).

40. Damiani, C. et al. Paternal transmission of symbiotic bacteria in malaria vectors. Curr. Biol. 18, R1087–R1088 (2008).

41. Whitten, M. M. A. et al. Symbiont-mediated RNA interference in insects. Proc. Biol. Sci. 283, 20160042 (2016).

42. Pontes, M. H. & Dale, C. Lambda red-mediated genetic modification of the insect endosymbiont *Sodalis glossinidius*. Appl. Environ. Microbiol. 77, 1918–1920 (2011).

43. Kim, J. K. et al. Bacterial Cell Wall Synthesis Gene *uppP* Is Required for *Burkholderia* Colonization of the Stinkbug Gut. Appl. Environ. Microbiol. 79, 4879–4886 (2013).

44. Schmidt, K. et al. Integration host factor regulates colonization factors in the bee gut symbiont *Frischella perrara*. eLife 12, e76182 (2023).

45. van Aartsen, J. J. & Rajakumar, K. An optimized method for suicide vector-based allelic exchange in *Klebsiella pneumoniae*. J. Microbiol. Methods 86, 313–319 (2011).

46. Motta, E. V. S., Powell, J. E., Leonard, S. P. & Moran, N. A. Prospects for probiotics in social bees. Philos. Trans. R. Soc. Lond. B. Biol. Sci. 377, 20210156 (2022).

47. Chhun, A. et al. Engineering a symbiont as a biosensor for the honey bee gut environment. 2023.02.02.526826 Preprint at 10.1101/2023.02.02.526826 (2023).

48. Baba, T. et al. Construction of *Escherichia coli* K-12 in-frame, single-gene knockout mutants: the Keio collection. Mol. Syst. Biol. 2, 2006.0008 (2006).

49. Murphy, K. C. λ Recombination and Recombineering. EcoSal Plus 7, 10.1128/ecosalplus.ESP-0011-2015 (2016).

50. Mosberg, J. A., Lajoie, M. J. & Church, G. M. Lambda Red Recombineering in *Escherichia coli* Occurs Through a Fully Single-Stranded Intermediate. Genetics 186, 791–799 (2010).

51. Reyrat, J. M., Pelicic, V., Gicquel, B. & Rappuoli, R. Counterselectable markers: Untapped tools for bacterial genetics and pathogenesis. Infect. Immun. 66, 4011–4017 (1998).

52. Benchling [Biology Software]. (2023).

53. Lee, M. E., DeLoache, W. C., Cervantes, B. & Dueber, J. E. A Highly Characterized Yeast Toolkit for Modular, Multipart Assembly. ACS Synth. Biol. 4, 975–986 (2015).

54. Wick, R. R., Judd, L. M., Gorrie, C. L. & Holt, K. E. Completing bacterial genome assemblies with multiplex MinION sequencing. *Microb*. Genomics 3, e000132 (2017).

55. Chen, S., Zhou, Y., Chen, Y. & Gu, J. fastp: an ultra-fast all-in-one FASTQ preprocessor. Bioinforma. Oxf. Engl. 34, i884–i890 (2018).

56. Deatherage, D. E. & Barrick, J. E. Identification of mutations in laboratory-evolved microbes from next-generation sequencing data using breseq. Methods Mol. Biol. 1151, 165–188 (2014).

57. Stead, C. M., Zhao, J., Raetz, C. R. H. & Trent, M. S. Removal of the outer Kdo from *Helicobacter pylori* lipopolysaccharide and its impact on the bacterial surface. Mol. Microbiol. 78, 837–852 (2010).

58. Strack, R. L. et al. A Rapidly Maturing Far-Red Derivative of DsRed-Express2 for Whole-Cell Labeling. Biochemistry 48, 8279–8281 (2009).

