## Supplementary material for "Single-step genome engineering in the bee gut symbiont *Snodgrassella alvi*": Table S3

**Predicted mutations**

| position | mutation | PL165 | PL224 | PL230 | PL236 | PL242 | PL246 | PL271 | AZA75 | AZA76 | AZA77 | annotation | gene | description |
|---|---|---|---|---|---|---|---|---|---|---|---|---|---|---|
| 859,190 | T→C | | | | | | | 100% | | | | F604L (TTT→CTT) | *aceE* → | pyruvate dehydrogenase (acetyl-transferring), homodimeric type |
| 1,009,711 | C→A | | | | | | | | 100% | 100% | 100% | S336Y (TCC→TAC) | *SALWKB2_RS11610* → | type VI secretion system Vgr family protein |
| 1,092,450 | C→T | | 100% | 100% | 100% | 100% | 100% | 100% | 100% | 100% | 100% | P155L (CCA→CTA) | *SALWKB2_RS05015* → | lysophospholipid acyltransferase family protein |
| 1,527,570 | Δ69 bp | 100% | | | | | | | | | | coding (129-197/216 nt) | *SALWKB2_RS07000* → | hypothetical protein |
| 1,603,327 | Δ1 bp | | 100% | | | | | | | | | intergenic (+83/-80) | *SALWKB2_RS07325* → / → *SALWKB2_RS07330* | DUF1415 domain-containing protein/sodium:proton antiporter |
| 1,603,364 | Δ1 bp | | | | 100% | | 100% | | | | | intergenic (+120/-43) | *SALWKB2_RS07325* → / → *SALWKB2_RS07330* | DUF1415 domain-containing protein/sodium:proton antiporter |
| 1,603,369 | Δ16 bp | 100% | | | | | | | | | | intergenic (+125/-23) | *SALWKB2_RS07325* → / → *SALWKB2_RS07330* | DUF1415 domain-containing protein/sodium:proton antiporter |
| 1,603,376 | Δ7 bp | Δ | | | | | | | 100% | 100% | 100% | intergenic (+132/-25) | *SALWKB2_RS07325* → / → *SALWKB2_RS07330* | DUF1415 domain-containing protein/sodium:proton antiporter |
| 1,648,618 | G→T | | 100% | | | | | | | | | G248* (GGA→TGA) | *SALWKB2_RS07515* → | autotransporter outer membrane beta-barrel domain-containing protein |
| 1,649,272 | Δ20 bp | | | 100% | | | | | | | | coding (1396-1415/5916 nt) | *SALWKB2_RS07515* → | autotransporter outer membrane beta-barrel domain-containing protein |
| 1,649,716 | +T | | | | 100% | | | | | | | coding (1840/5916 nt) | *SALWKB2_RS07515* → | autotransporter outer membrane beta-barrel domain-containing protein |
| 1,650,595 | Δ1 bp | | | | | | 100% | | | | | coding (2719/5916 nt) | *SALWKB2_RS07515* → | autotransporter outer membrane beta-barrel domain-containing protein |
| 1,855,359 | $(T)_{8\rightarrow7}$ | | 100% | 100% | 100% | 100% | 100% | | | | | coding (1120/1575 nt) | *SALWKB2_RS08395* ← | HAMP domain-containing sensor histidine kinase |
| 1,914,213 | A→C | 100% | | | | | | | | | | N163K (AAT→AAG) | *SALWKB2_RS08660* ← | response regulator transcription factor |
